# Evolutionarily Conserved and Divergent Mechanisms of Dual Ca^2+^ Sensors in Synaptic Vesicle Exocytosis

**DOI:** 10.1101/2025.09.30.679486

**Authors:** Lei Li, Jingyao Xia, Jiafan Wang, Xiaochun Yu, Jiayi Hu, Janet Richmond, Haowen Liu, Zhitao Hu

**Author notes:** These authors contributed equally to this work (Lei Li, Jingyao Xia). Lead Contact: Zhitao Hu.

## Abstract

Neurotransmitter release at the *C. elegans* neuromuscular junction is governed by a dual Ca²⁺ sensor system composed of SNT-1 and SNT-3, which are functional analogs of synaptotagmin-1 and -7 (Syt1/Syt7) in mammalian central synapses. In this study, we investigated how SNT-1 and SNT-3 mediate fast and slow neurotransmitter release through their potential interactions with the SNARE complex and their polybasic motifs. AlphaFold 3 models of SNT-1–SNARE and SNT-3–SNARE complexes accurately recapitulated the canonical Syt1 C2B–SNARE primary interface (Zhou et al., 2015, *Nature*) and precisely identified conserved binding residues within the C2B domains, as well as in SNAP-25 and Syntaxin, highlighting the evolutionary conservation of this interaction. Electrophysiological analyses using targeted mutagenesis demonstrated that both SNT-1 and SNT-3 require C2B–SNARE interactions and polybasic motifs within their C2 domains to drive evoked fast and slow neurotransmitter release. Notably, SNT-1 and SNT-3 exhibited differential dependence on distinct regions of the C2B–SNARE interface and their respective polybasic motifs, suggesting that Ca²⁺-triggered fast and slow release operate via distinct mechanistic strategies. Furthermore, we found that SNT-1 mediates spontaneous neurotransmitter release through multiple pathways, involving not only the primary C2B–SNARE interface but also additional putative SNARE-binding interactions. Together, our findings uncover both conserved and divergent mechanisms for synaptic exocytosis regulated by the dual Ca^2+^ sensors in *C. elegans*.

## Introduction

Neurotransmitters are released from presynaptic nerve terminals through synaptic vesicle (SV) exocytosis—a process regulated by the SNARE (soluble N-ethylmaleimide sensitive factor attachment protein receptor) complex, the core component of the membrane fusion machinery (Chapman, 2008; Jahn and Scheller, 2006; Sudhof, 2012). This process is tightly orchestrated by SNARE-binding proteins, such as Munc13, Munc18, complexin, and synaptotagmin (Sudhof and Rizo, 2011). Among these, Syt1, a SV protein, has been established as the primary Ca²⁺ sensor responsible for fast and synchronous neurotransmitter release in different model organisms (Geppert et al., 1994; Lee et al., 2013; Li et al., 2018a; Sudhof, 2013; Yoshihara and Littleton, 2002). Other synaptotagmin isoforms, such as Syt2 and Syt9, also function as fast Ca²⁺ sensors in specific neuron groups (Kochubey et al., 2016; Xu et al., 2007). The typical architecture of these Syts includes a short N-terminal luminal segment, a transmembrane domain that anchor the protein on either SVs or the plasma membrane, an unstructured linker, and the tandem C2 domains—C2A and C2B—both of which bind Ca²⁺ via conserved aspartate residues in their Ca²⁺-binding loops (Chapman, 2008). Ca²⁺ binding, particularly to the C2B domain of Syt1, has been shown to be essential for Syt1 to trigger SV fusion (Li et al., 2021; Li et al., 2018a; Littleton et al., 2001; Shin et al., 2003).

Recent work has identified Syt7 as a secondary Ca²⁺ sensor that mediates slow, asynchronous neurotransmitter release at central synapses. Unlike Syt1, Syt7 knockout does not impair basal synaptic transmission but abolishes the residual asynchronous release observed in Syt1-deficient neurons (Bacaj et al., 2013). Moreover, SV priming is nearly eliminated in Syt1/Syt7 double knockout neurons, whereas it remains intact in either single knockout, suggesting that Syt1 and Syt7 have partially redundant roles in SV priming (Bacaj et al., 2015). Together, these findings support a dual Ca²⁺ sensor model in which Syt1 and Syt7 act as fast and slow Ca²⁺ sensors, respectively, to control distinct temporal components of synaptic transmission. In addition to driving asynchronous release, Syt7 also plays roles in synaptic facilitation and vesicle pool replenishment (Chen et al., 2017; Jackman et al., 2016), and contributes to frequency invariance in certain depressing synapses (Turecek et al., 2017).

Despite two decades of research, the precise molecular mechanisms by which Syt1 and Syt7 regulate SV exocytosis remain incompletely understood. Biochemical and functional studies have revealed that Syt1’s C2 domains interact with the SNARE complex (Chapman et al., 1995; Chapman and Jahn, 1994; Sutton et al., 1995; Wang et al., 2016; Zhang et al., 2002), a critical step for SV fusion. Although the structures of isolated Syt1 C2 domains and the SNARE complex have been solved, a detailed structural model of the full Syt1–SNARE complex remained elusive for many years. Recently, work from Brunger’s group provided the first crystal structure of the Syt1–SNARE complex at atomic-resolution levels (Zhou et al., 2015), revealing multiple contact interfaces between the C2 domains of Syt1 and the SNARE proteins. Moreover, mutations of the residues involved in the primary interface of Syt1-SNARE significantly impaired the function of Syt1 as the fast Ca^2+^ sensor. These findings offer new mechanistic insights into synaptotagmin-mediated synaptic exocytosis.

Interaction of Syt1 C2 domains with the plasma membrane also plays critical roles in triggering SV exocytosis. The C2 domains respond to increases in cytoplasmic Ca^2+^ levels, which prompts side chains of the Ca^2+^-binding loops to quickly insert into lipid membranes containing negatively charged phospholipids (Bai et al., 2000; Bai et al., 2004; Bradberry et al., 2019; Chapman and Davis, 1998; Grushin et al., 2019). It’s well-established that this membrane insertion by Syt1 speeds up the SNARE-driven SV fusion process, both *in vitro* and in cultured neurons (Bai et al., 2016). In addition to the Ca^2+^ binding loops, the polybasic motifs in both C2 domains have also been shown to interact with the plasma membrane. The four consecutive lysine residues in each domain form a polylysine patch which has been shown to bind to PIP2 lipid clusters on the plasma membrane, in a Ca^2+^-independent manner (Bai et al., 2004; Bradberry et al., 2019; Loewen et al., 2006; Park et al., 2015). It has been proposed that the SNARE-interaction and the membrane-interaction in Syt1 occur simultaneously before exocytosis, and Ca^2+^ influx promotes these interactions and remodels the membrane to promote SV fusion (Zhou et al., 2015).

Our previous studies have revealed a dual Ca²⁺ sensor system in *C. elegans*, consisting of SNT-1 (the worm homolog of Syt1) and SNT-3, both of which share high sequence similarity with Syt1 (Li et al., 2021). Unlike most other synaptotagmin isoforms in *C. elegans* and mammals, SNT-3 uniquely lacks a transmembrane domain. Our functional and biochemical studies have demonstrated that SNT-1 and SNT-3 act as fast and slow Ca²⁺ sensors triggering evoked fast and slow release (Li et al., 2021). Remarkably, the SNT-1/SNT-3 system in *C. elegans* mirrors the mammalian Syt1/Syt7 system in several functional aspects. First, the primary (SNT-1, Syt1) and secondary (SNT-3, Syt7) Ca²⁺ sensors exhibit distinct subcellular localization (Brose et al., 1992; Li et al., 2021; Li et al., 2018b; Littleton et al., 1993; Maximov et al., 2008; Sugita et al., 2001). Second, loss of the primary sensor leads to severe impairment of fast synaptic transmission, whereas deletion of the secondary sensor alone does not affect basal transmission but abolishes the residual release in the absence of the primary sensor (Bacaj et al., 2013; Lee et al., 2013; Li et al., 2021; Li et al., 2018b; Maximov et al., 2008; Nishiki and Augustine, 2004b; Weber et al., 2014; Xu et al., 2007; Yoshihara and Littleton, 2002). Third, primary and secondary Ca²⁺ sensors redundantly contribute to SV priming (Bacaj et al., 2015; Li et al., 2021). Together, these shared features establish *C. elegans* as a powerful model for dissecting evolutionarily conserved mechanisms of dual Ca²⁺ sensor–mediated synaptic transmission.

Despite the work done on the dual Ca²⁺ sensors, the precise mechanisms by which they trigger SV exocytosis remain unclear. Do they engage SNAREs and the plasma membrane similarly to Syt1? Are these interactions essential for their roles as Ca²⁺ sensors? What underlies their distinct functional contributions in fast and slow neurotransmitter release? In this study, we investigated the roles of conserved residues in SNT-1 and SNT-3 that correspond to the SNARE-interacting surfaces identified in the Syt1–SNARE complex. Our findings demonstrate that the primary C2B–SNARE interface identified in the Syt1– SNARE structure is essential for the function of both Ca²⁺ sensors in *C. elegans*. However, the relative contributions of specific subregions within this interface differ between SNT-1 and SNT-3. Moreover, the polybasic motifs in the C2 domains of SNT-1 and SNT-3 also play critical roles in SV exocytosis, though their contributions differ between the C2A and C2B domains of these sensors. Additionally, we found that SNT-1 participates in multiple potential modes of SNARE interaction, which collectively support both Ca²⁺-dependent and Ca²⁺-independent spontaneous neurotransmitter release. Altogether, our findings suggest that the dual Ca²⁺ sensors in *C. elegans* utilize both conserved and divergent molecular mechanisms to trigger synaptic exocytosis, providing important insight into the evolution of synaptic transmission mechanisms.

## Results

### Mammalian Syt1 restored behavioral and transmission defects in *snt-1* mutant

To investigate whether *C. elegans* SNT-1 and mammalian Syt1 share conserved functions in the nervous system, we expressed rat Syt1 cDNA in the nervous system of the worm *snt-1* mutants. Our results showed that both SNT-1 and Syt1 expression significantly rescued the reduced body length observed in *snt-1* mutants (Figure 1A, B). Additionally, Syt1 restored the thrashing rate defect in the *snt-1* mutants to a level comparable to that of wild-type SNT-1 (Figure 1A, B). These findings suggest that rat Syt1 functions effectively in worm neurons.

**Figure 1.**
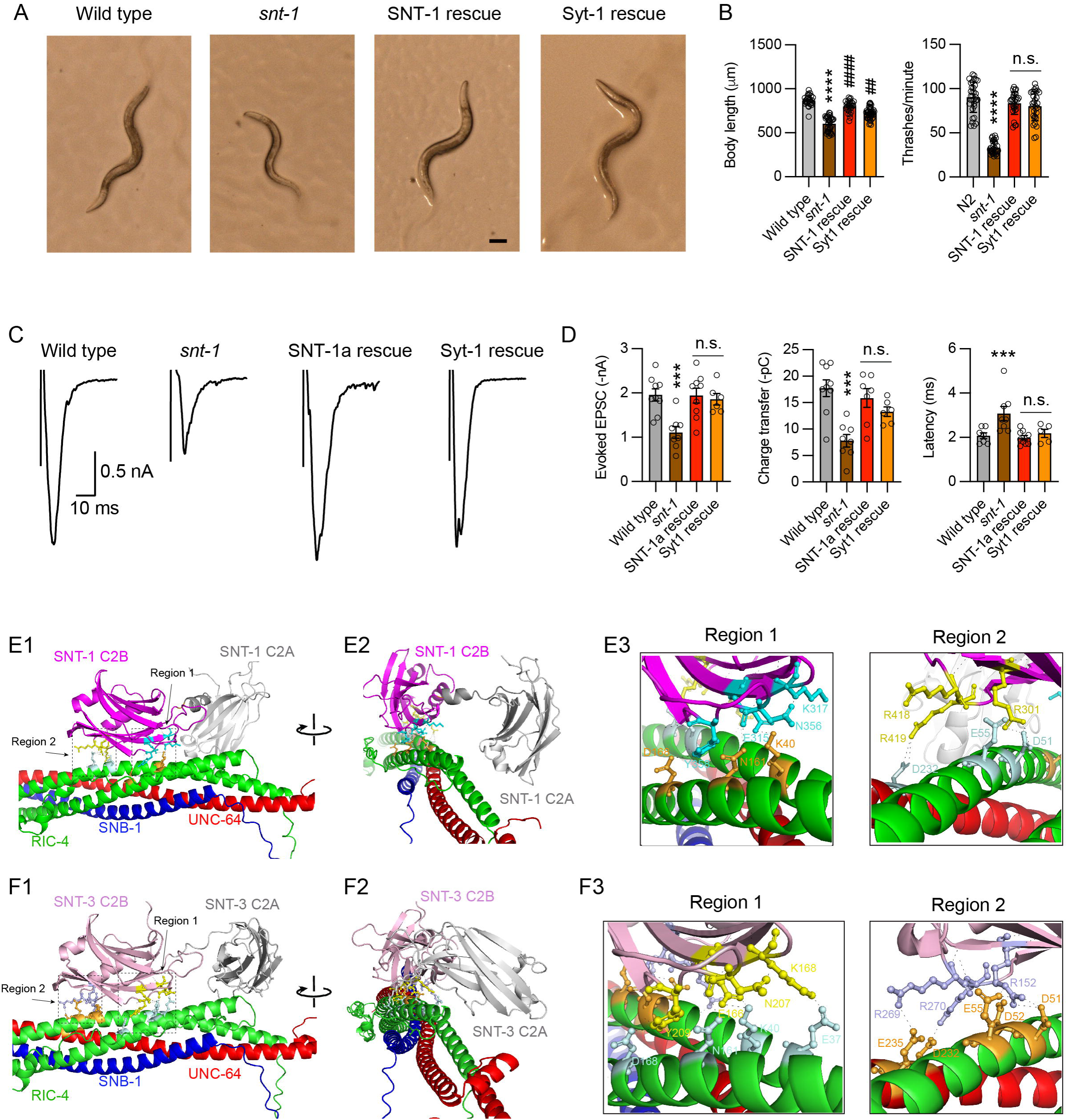
SNT-1 and SNT-3 exhibit conserved interactions with the SNARE complex similar to Syt1. **(A)** Representative images of wild-type, *snt-1* mutant, SNT-1 rescue, and Syt1 rescue worms. Both SNT-1 and Syt1 expression rescued the body size defect observed in *snt-1* mutants. **(B)** Quantification of body length and thrash corresponding to the genotypes shown in (A). Data are presented as mean ± SEM (**** *p* < 0.0001 vs. wild-type; ## *p* < 0.01, #### *p* < 0.0001 vs. *snt-1* mutant; n.s., not significant compared to SNT-1 rescue; one-way ANOVA). **(C)** Representative traces of stimulus-evoked EPSCs recorded from wild-type, *snt-1* mutant, SNT-1 rescue, and Syt1 rescue animals under 1 mM Ca²⁺ conditions. **(D)** Quantification of evoked EPSC amplitude, charge transfer, and latency. Data are presented as mean ± SEM (*** *p* < 0.001 vs. wild-type; n.s., not significant compared to SNT-1 rescue; one-way ANOVA). **(E1, E2)** AlphaFold 3 models of the SNT-1–SNARE complex, shown in front view (E1) and side view (E2). Two binding regions at the interface are highlighted by dashed squares. **(E3, E4)** Key residues involved in the SNT-1–SNARE primary interface: region 1 (E3; E315/K317/N356/Y358 in SNT-1 and K40/N161/D168 in RIC-4 (SNAP-25)) and region 2 (E4; R301/R418/R419 in SNT-1, D51/E55 in RIC-4, and D232 in UNC-64 (syntaxin)). **(F1, F2)** AlphaFold 3 models of the SNT-3–SNARE complex, shown in front view (F1) and side view (F2). **(F3, F4)** Key residues involved in the SNT-3–SNARE primary interface: region 1 (F3; E166/K168/N207/Y209 in SNT-3 and E37/K40/N161/D168 in RIC-4) and region 2 (F4; R152/R269/R270 in SNT-3, D51/D52/E55 in RIC-4, and D232/E235 in UNC-64).

To directly assess synaptic transmission, we recorded stimulus-evoked <u>e</u>xcitatory <u>p</u>ost<u>s</u>ynaptic <u>c</u>urrents (evoked EPSCs) at *C. elegans* neuromuscular junctions (NMJs) and found that Syt1 expression robustly restored evoked EPSCs to levels similar to SNT-1-rescued animals (Figure 1C, D). Additionally, the increased latency of evoked EPSCs observed in *snt-1* mutants was effectively rescued by Syt1, suggesting that Syt1 functions as a fast Ca^2+^ sensor in worms, triggering fast neurotransmitter release through mechanisms likely analogous to those in mammals. Furthermore, the frequencies of miniature excitatory and inhibitory postsynaptic currents (mEPSCs and mIPSCs, referred to as spontaneous release or minis), which were markedly reduced in *snt-1* mutants, were also significantly rescued by Syt1 (Figure 1-Supplemental 1). These findings demonstrate a high degree of functional conservation between *C. elegans* SNT-1 and mammalian Syt1 in mediating Ca²⁺- dependent neurotransmitter release.

### SNT-1 possesses conserved SNARE-binding sites

The ability of Syt1 to rescue behavior and evoked EPSCs in *snt-1* mutants suggests that SNT-1 and Syt1 may share similar molecular mechanisms of action, particularly in their interactions with the SNARE complex. However, the absence of a crystal structure for the SNT-1–SNARE complex limits our understanding of how SNT-1 binds SNAREs in *C. elegans*. Given the high predictive accuracy of AlphaFold 3—demonstrated by its near-perfect prediction of the Munc13-1 structure compared to cryo-EM data (Grushin et al., 2022)—we asked whether AlphaFold 3 could also be used to predict the structural interface between SNT-1 and the SNARE complex.

As a proof of concept, we first tested the ability of AlphaFold 3 to recapitulate the known Syt1–SNARE interaction. As expected, AlphaFold 3 successfully predicted a stable interaction between the Syt1 C2B domain and the SNARE complex. The predicted C2B–SNARE interface closely resembled the primary binding interface observed in crystallographic studies of the Syt1-SNARE complex, with two distinct binding regions (region 1 and region 2) located at opposite ends of the C2B domain (Figure 1-Supplemental 2A). Moreover, AlphaFold 3 precisely identified most of the key binding residues in both regions of C2B, as well as in SNAP25 (Figure 1-Supplemental 2B, C). The only notable deviation from the crystal structure was the absence of the predicted interaction between C2B region 2 residues R398 and R399 with Syntaxin (Figure 1-Supplemental 2D). In very few models, the R399 residue displayed interaction with the D231 residue of syntaxin. Nevertheless, this predicted structural arrangement supports a stable, high-affinity interaction and validates the use of AlphaFold 3 for modeling SNARE-binding interfaces.

Using AlphaFold 3, we found that the C2B domain of SNT-1 and the *C. elegans* SNARE proteins (UNC-64, RIC-4, and SNB-1) form a predicted interface highly similar to the Syt1 C2B–SNARE interaction. The model also revealed two primary binding regions located at opposite ends of the C2B domain, mediated by conserved residues in both SNT-1 and RIC-4 (the worm homolog of SNAP-25) (Figure 1E1-E3). Consistent with the Syt1– SNARE model, AlphaFold 3 did not consistently predict the interaction between SNT-1 C2B residues R398 and R399 and UNC-64 (the worm Syntaxin-1A). Although not all residues from the crystallographically defined primary interface were reproduced, several key residues—E295, N336, and Y338 in region 1, and R281 in region 2—were consistently identified in nearly all AlphaFold 3 predictions for both Syt1– and SNT-1–SNARE complexes. This conservation strongly suggests thats SNT-1 interacts with the SNARE complex in worms through a mechanism analogous to Syt1 in mammals, mediating fast neurotransmitter release.

### SNT-3 displays similar SNARE interactions as Syt1 and SNT-1

The high sequence similarity between the C2B domains of SNT-1 and SNT-3, along with the functional requirement of the SNT-3 C2B domain for evoked EPSCs (Li et al., 2021), led us to hypothesize that this secondary Ca^2+^ sensor also promotes synaptic exocytosis by interacting with SNAREs. Supporting this, AlphaFold 3 predicted a similar two-region interface between SNT-3 C2B and the SNARE complex. Notably, nearly all residues implicated in the Syt1 C2B–SNARE interface were accurately identified in the SNT-3 model (Figure 1F1-F3, Figure 1-Supplemental 2D). In particular, residues R269 and R270 in region 2 were predicted to interact with UNC-64 in the majority of models, suggesting that this interface is also conserved in SNT-3. These observations strongly support the hypothesis that SNT-3 engages SNARE proteins through a conserved mechanism to mediate evoked slow neurotransmitter release.

### The C2B–SNARE interface is essential for evoked fast and slow neurotransmitter release

To assess the functional importance of the predicted C2B–SNARE interface in SNT-1 and SNT-3, we generated quintuple point mutations targeting five conserved residues implicated in SNARE binding, analogous to those identified in Syt1. These mutations included E315A, Y358W, R301A, R418A, and R419A in SNT-1 (hereafter referred to as SNT-1^Quintuple^), and E166A, Y209W, R152A, R269A, and R270A in SNT-3 (SNT-3^Quintuple^). The mutant constructs were expressed in the nervous system of *snt-1* and *snt-1; snt-3* double mutants, respectively (note that the effects of *snt-3* knockout are detectable only in the *snt-1* mutant background, similar to Syt7 knockout). In contrast to their wild-type counterparts, neither SNT-1^Quintuple^ nor SNT-3^Quintuple^ rescued evoked EPSCs in the respective mutant backgrounds (Figure 2A-D). Moreover, the quintuple mutations in SNT-1 resulted in significantly prolonged EPSC latency, comparable to that observed in *snt-1* mutants. These defects are unlikely to result from reduced expression, as prior studies in *C. elegans* SNT-1 and *Drosophila* Syt1 have shown that deletion of the entire C2B or C2A domain does not compromise protein expression (Lee et al., 2013; Li et al., 2018a). These results demonstrate that conserved residues in SNT-1 and SNT-3 are critical for SNARE complex interactions, and that the potential C2B-SNARE interface is essential for mediating both fast and slow modes of evoked neurotransmitter release.

**Figure 2.**
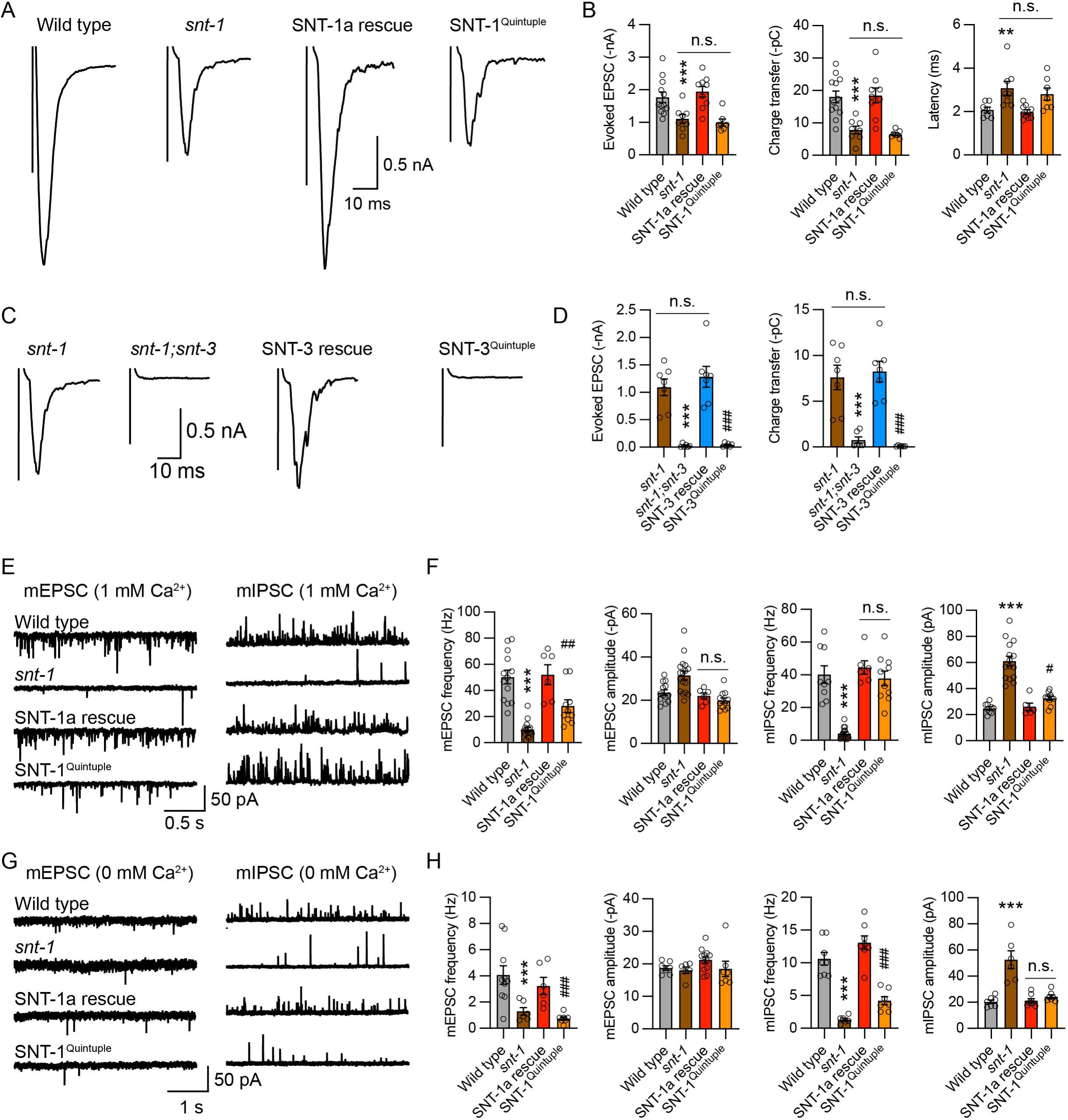
The conserved SNARE-binding residues are critical for synaptic transmission mediated by SNT-1 and SNT-3. **(A)** Representative traces of evoked EPSCs recorded from wild-type, *snt-1* mutant, SNT-1 rescue, and SNT-1^Quintuple^ rescue animals. **(B)** Quantification of evoked EPSC amplitude, charge transfer, and latency from the same genotypes shown in A. The quintuple mutations in SNT-1 abolished evoked fast neurotransmitter release. **(C)** Representative traces of evoked EPSCs from *snt-1* mutant, *snt-1;snt-3* double mutant, SNT-3 rescue, and SNT-3^Quintuple^ rescue animals. Wild-type SNT-3 and SNT-3^Quintuple^ were expressed in the *snt-1;snt-3* double mutant background. **(D)** Quantification of evoked EPSCs in the genotypes shown in C. Similar to SNT-1, the quintuple mutations in SNT-3 abolished evoked slow neurotransmitter release. **(E, F)** Example traces and quantification of mEPSC and mIPSC frequencies and amplitudes (recorded in 1 mM Ca²⁺) for the genotypes in A. The quintuple mutations in SNT-1 reduced mEPSC frequency by ∼50% without affecting mIPSC frequency. **(G, H)** Representative traces and quantification of mEPSCs and mIPSCs recorded under 0 mM Ca²⁺ conditions. In contrast to the 1 mM Ca²⁺ condition, the quintuple mutations strongly reduced both mEPSC and mIPSC frequencies. Despite this reduction, mIPSC amplitude remained unchanged. Data are presented as mean ± SEM (** *p* < 0.01, *** *p* < 0.001 vs. wild-type; # *p* < 0.05; ## *p* < 0.01; ### *p* < 0.001 vs. SNT-1 rescue or SNT-3 rescue; n.s., not significant; one-way ANOVA).

### The C2B–SNARE interface plays a modest role in Ca²⁺-dependent spontaneous release and a major role in Ca²⁺-independent conditions

Previous studies have shown that SNT-1 is essential for spontaneous neurotransmitter release, as *snt-1* mutants exhibit significantly reduced frequencies of mEPSCs and mIPSCs (Li et al., 2021; Li et al., 2018a). Interestingly, the mechanisms governing spontaneous and evoked release appear to differ: mutations that disrupt Ca²⁺ binding in the C2B domain abolish evoked EPSCs but leave spontaneous release largely unaffected (Li et al., 2018a). To examine whether the C2B–SNARE interface contributes to spontaneous release, we compared mEPSCs and mIPSCs in animals rescued with wild-type SNT-1 versus SNT-1^Quintuple^ in the presence of 1 mM Ca²⁺. The SNT-1^Quintuple^ mutant exhibited a ∼50% reduction in mEPSC frequency, while mIPSCs remained unchanged, suggesting that the C2B–SNARE interaction contributes modestly to Ca²⁺-triggered spontaneous acetylcholine (ACh) release.

Spontaneous release also occurs in the absence of extracellular Ca²⁺ at the *C. elegans* NMJ, and this Ca^2+^-independent release is abolished in mutants lacking the core SNARE components, including UNC-64, SNB-1, and RIC-4 (Liu et al., 2018). Given that the C2B–SNARE interaction can occur independently of Ca²⁺ (Zhou et al., 2015), we next examined the impact of the SNT-1 quintuple mutations under 0 mM Ca²⁺ conditions. Remarkably, both mEPSC and mIPSC frequencies were dramatically reduced in SNT-1^Quintuple^ rescue animals under these conditions, indicating that the C2B–SNARE interface is critical for Ca²⁺- independent spontaneous release. Notably, mIPSC frequency in the SNT-1^Quintuple^ background remained higher than in *snt-1* null mutants (Figure 2H). These results, together with the results of spontaneous release in the presence of Ca^2+^, suggest the presence of additional mechanisms in SNT-1-mediated spontaneous release.

To investigate the Ca²⁺-independence of the SNT-1-SNARE interaction, we hypothesized that disrupting the Ca²⁺-binding capability of SNT-1 would not affect spontaneous release in the absence of Ca²⁺. To test this, we introduced mutations by replacing the third and fourth aspartate residues with asparagines in the C2A domain, C2B domain, or both (Figure 1, Supplemental-3; termed SNT-1^C2A-D3,4N^, SNT-1^C2B-D3,4N^, and SNT-1^C2AB-D3,4N^), and assessed their rescue ability in *snt-1* mutant strains. As expected, both mEPSCs and mIPSCs in the mutant strains were restored to levels comparable to those of the wild-type SNT-1 in 0 mM Ca²⁺ conditions (Figure 2, Supplemental-1A-D). These results indicate that the SNARE interaction of SNT-1 is independent of its Ca²⁺-binding ability.

In the presence of Ca²⁺, the Ca²⁺-binding mutations in either the C2A or C2B domains did not significantly affect mEPSCs or mIPSCs, mirroring the results observed in 0 mM Ca²⁺ (Figure 2, Supplemental-1E-H). While mEPSCs were reduced by approximately 40% by the Ca²⁺- binding mutations in both C2 domains, mIPSCs remained unaffected. These findings are consistent with those resulting from C2B SNARE-binding mutations (Figure 2E, F), suggesting that simultaneous disruption of Ca²⁺ binding in the C2 domains may influence their conformation and relative positioning, thereby affecting the C2B-SNARE interaction.

In addition to frequency changes, *snt-1* mutants also exhibited increased mEPSC and mIPSC amplitudes under both Ca²⁺-containing and Ca²⁺-free conditions (Figure 2F, H). Our prior electron microscopy analysis revealed significantly enlarged SVs in *snt-1* mutants, with increased average SV diameter (Li et al., 2021). This SV enlargement is likely a consequence of impaired endocytosis in the absence of functional SNT-1. Supporting this idea, enlarged SV size and increased miniature amplitudes have also been observed in other mutants with established endocytic defects (Mullen et al., 2012) (Hu lab, unpublished data). Interestingly, the only difference from this pattern was observed in mEPSC amplitude under 0 mM Ca²⁺ conditions, which was not increased in *snt-1* mutants. This anomaly is likely due to a postsynaptic regulation. Our previous work has shown that direct application of ACh onto body wall muscles evoked smaller currents in 0 mM Ca²⁺ recordings compared to 1mM Ca^2+^ conditions, whereas GABA-induced currents remained unchanged (Liu et al., 2018), suggesting a specific reduction in postsynaptic cholinergic sensitivity under these conditions. Notably, although the SNT-1 quintuple mutations caused a marked reduction in mIPSC frequency, they did not increase mIPSC amplitude as observed in *snt-1* mutants (Figure 2H). One possible explanation is that the C2B–SNARE interface is primarily involved in the exocytosis process but not in endocytosis. Thus, SNT-1^Quintuple^ may retain normal function in SV endocytosis.

### The two binding regions of the C2B-SNARE primary interface play distinct roles in SNT-1 and SNT-3 function

The primary interface between the C2B domain and the SNARE complex comprises two conserved binding regions located at opposite ends of the C2B domain (Figure 1A1, B1), forming a stable interaction critical for the functional activity of SNT-1 and SNT-3 (Figure 2). To dissect the individual contributions of these regions, we introduced point mutations into conserved EY residues in region 1 (E315A and Y358W in SNT-1; E166A and Y209W in SNT-3) and RR residues in region 2 (R418 and R419 in SNT-1; R269 and R270 in SNT-3), and examined their effects on SNT-1- and SNT-3-mediated SV release.

In SNT-1, either E315A/Y358W or R418A/R419A mutations completely abolished SNT-1-dependent evoked EPSCs (Figure 3A, B), indicating that both regions are indispensable for its role in mediating fast neurotransmitter release. By contrast, in the *snt-1;snt-3* double mutant background, SNT-3-mediated evoked EPSCs were eliminated by the R269A/R270A mutations but were unaffected by the E166A/Y209W mutations (Figure 3C, D). These results suggest that while both regions are essential for SNT-1 function, only region 2 is critical for SNT-3-mediated slow release, revealing mechanistic divergence in how the two Ca^2+^ sensors engage the C2B–SNARE interface. Since region 1 includes two additional binding residues (K297 and N336 in Syt1, and their conserved counterparts in SNT-1 and SNT-3; see Figure 1 and Figure 1 supplemental 1), which may contribute to partial interface formation, the results of EY mutations likely reflect minimal binding requirements between the C2B domain and SNARE complex for triggering fast versus slow neurotransmitter release (see discussion).

**Figure 3.**
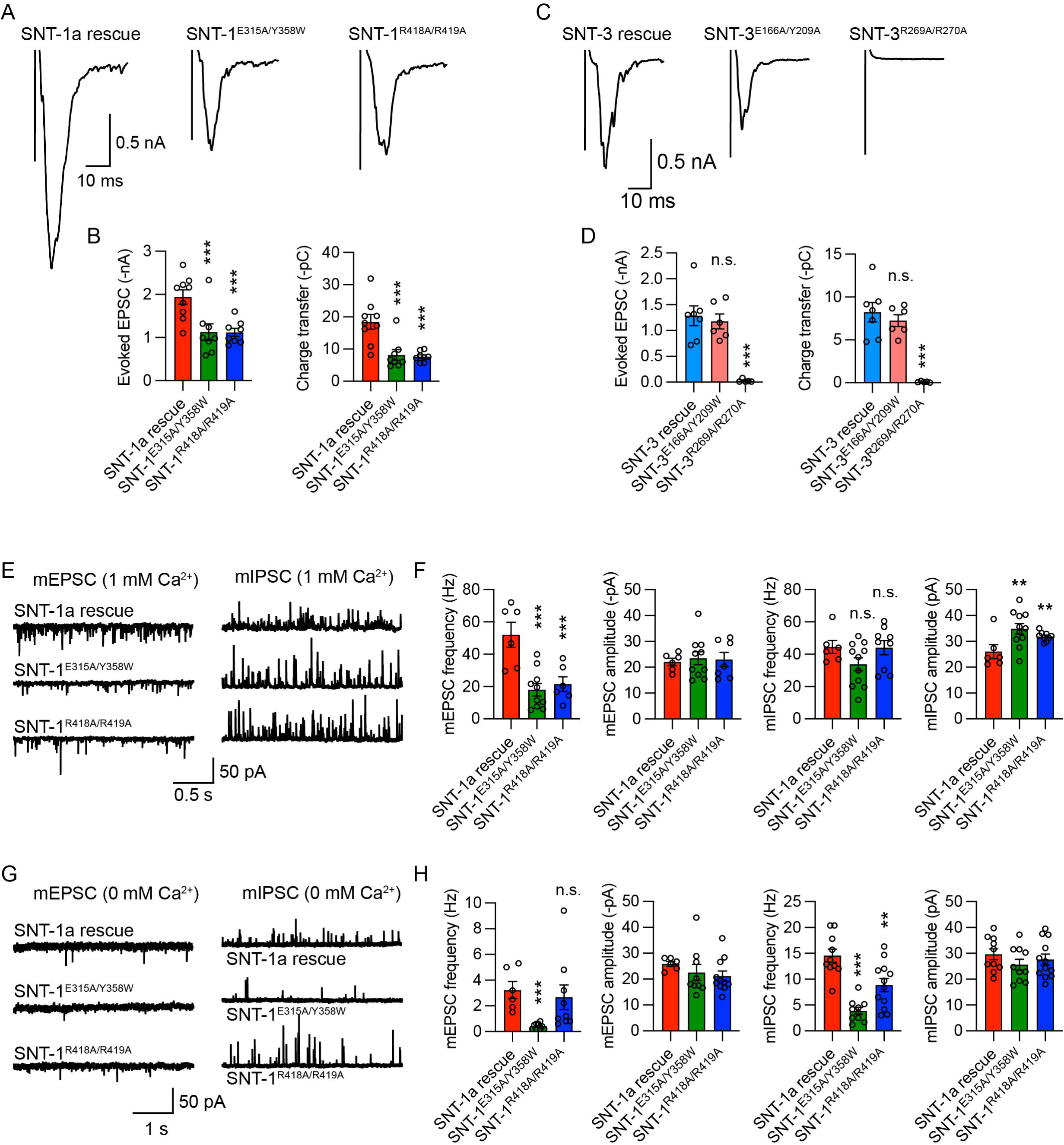
Distinct contributions of the two binding regions in SNT-1–SNARE and SNT-3–SNARE interface to synaptic transmission. **(A)** Representative traces of evoked EPSCs recorded from *snt-1* mutants rescued with wild-type SNT-1, SNT-1^E315A/Y358W^, and SNT-1^R418A/R419A^. **(B)** Averaged amplitude and charge transfer of evoked EPSCs shown in A. Both E315A/Y358W (region 1) and R418A/R419A (region 2) mutations completely abolished evoked fast release mediated by SNT-1. **(C)** Representative evoked EPSC traces from *snt-1;snt-3* double mutants rescued by wild-type SNT-3, SNT-3^E166A/Y209A^, and SNT-3^269A/R270A^. **(D)** Quantification of evoked EPSC amplitude and charge transfer in the genotypes shown in C. The R269A/R270A (region 2) mutations abolished evoked EPSCs mediated by SNT-3, while the E166A/Y209A (region 1) mutations had no effect. **(E, F)** Representative traces and quantification of mEPSC and mIPSC frequencies and amplitudes (recorded under 1 mM Ca²⁺) for the genotypes in A. Both E315A/Y358W and R418A/R419A mutations in SNT-1 reduced mEPSC frequency by ∼50%, but did not affect mIPSC frequency. **(G, H)** Representative traces and quantification of mEPSCs and mIPSCs recorded under 0 mM Ca²⁺ conditions. Unlike in 1 mM Ca²⁺, the R418A/R419A mutations had no effect on mEPSCs and only mild effects on mIPSCs, whereas the E166A/Y209A mutations strongly decreased both mEPSC and mIPSC frequencies. Data are presented as mean ± SEM (** *p* < 0.01, *** *p* < 0.001 vs. SNT-1 or SNT-3 rescue; n.s., not significant compared to SNT-1 or SNT-3 rescue; one-way ANOVA).

To explore the role of these binding regions in spontaneous release, we next examined mEPSCs and mIPSCs in animals expressing mutant SNT-1. Under 1 mM Ca²⁺ conditions, both E315A/Y358W and R418A/R419A mutations led to a ∼50% reduction in mEPSC frequency, with no significant change in mIPSC frequency (Figure 3E, F), recapitulating the effect observed in the quintuple mutant (Figure 2E, F). This indicates that both binding regions contribute comparably to Ca²⁺-dependent spontaneous ACh release.

Interestingly, under 0 mM Ca²⁺ conditions, the E315A/Y358W mutations—but not the R418A/R419A mutations—resulted in a profound reduction of both mEPSC and mIPSC frequencies (Figure 3G, H), nearly matching the effect of the full quintuple mutations. These findings suggest that region 1 plays a more dominant role in Ca²⁺-independent spontaneous release. One possible explanation is that region 1 forms a tighter interaction with SNAREs in the absence of Ca²⁺, while Ca²⁺ enhances the binding strength of both regions, equalizing their contributions during Ca²⁺-dependent release. This model is supported by prior studies showing that Ca²⁺ can potentiate the interaction of Syt1 C2 domains with the SNARE complex (Chapman et al., 1995; Dai et al., 2007; Huang and Cafiso, 2008; Voleti et al., 2020).

### SNT-1 triggers spontaneous release through additional mechanisms beyond the primary SNARE interface

Given that spontaneous neurotransmitter release persists in the SNT-1 quintuple mutant under 1mM Ca²⁺ conditions, we hypothesized that SNT-1 may engage additional mechanisms beyond the primary C2B–SNARE interface to mediate spontaneous SV fusion. Structural studies of the Syt1–SNARE complex have identified two additional contact points: a secondary interface between the C2B domain and SNARE (involving K288 of Syt1 and E55 of SNAP-25), and a tertiary interface between the C2A domain and SNARE (involving R199 and R233 of Syt1, and D218 of Syntaxin and D57 of Synaptobrevin (Figure 1 Supplemental 3) (Zhou et al., 2015). Notably, the C2 domains adopt distinct orientations in the secondary and tertiary interfaces compared to the primary interface, indicating that these additional interfaces occur when C2B does not form a primary interface. Importantly, the residues involved in the secondary and tertiary interfaces of the Syt1-SNARE complex are also highly conserved in *C. elegans* SNT-1 and SNT-3 (Figure 1-Supplemental 3).

Unlike the primary interface, however, most AlphaFold 3-predicted models did not recapitulate the secondary or tertiary interfaces of the Syt1–SNARE complex derived from crystal structures. Only a minority of models predicted the interaction between K288 of Syt1 and E55 of SNAP-25, consistent with the secondary interface observed in the Syt1 structure (Figure 4A). Similarly, in some AlphaFold 3 models of the SNT-1–SNARE complex, a comparable interaction was observed between K308 in the SNT-1 C2B domain and E55 of RIC-4 (SNAP-25), suggesting the potential conservation of the secondary interface in SNT-1. However, the orientation of the C2B domain in these predicted models differed from that in the experimentally resolved Syt1–SNARE complex. In fact, in these models, the secondary and primary interfaces were predicted to occur simultaneously—a configuration not observed in structural studies. This discrepancy likely stems from the stable and consistent primary C2B–SNARE interface that dominates across AlphaFold 3 models, potentially constraining the conformational flexibility required for the C2B or C2A domains to adopt alternative orientations necessary for forming secondary or tertiary interfaces. Since structural studies have shown that the secondary and tertiary interfaces in Syt1 are formed in conformations that are mutually exclusive with the primary interface, it is not surprising that AlphaFold 3, which tends to favor the most stable interaction state, fails to capture these dynamic transitions. Therefore, the AlphaFold 3 modeling cannot exclude the possibility that SNT-1 and SNT-3, like Syt1, are capable of engaging in alternative SNARE-binding modes through conformational rearrangements. Such alternative interfaces may underlie SNT-1’s residual ability to support spontaneous neurotransmitter release even when the primary C2B–SNARE interface is disrupted.

**Figure 4.**
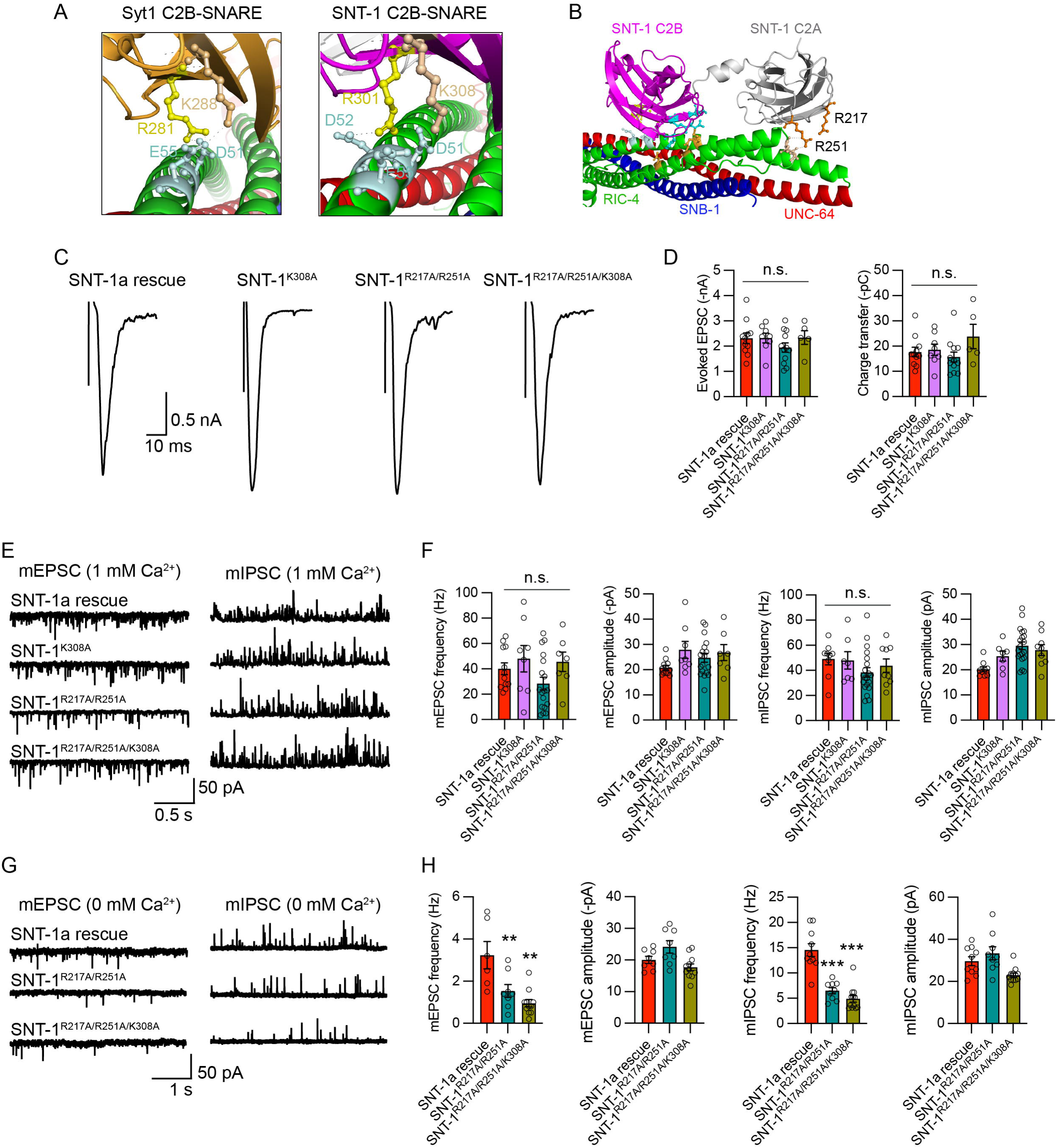
Residues mediating potential additional SNT-1–SNARE interactions are critical for Ca²⁺-independent release. **(A)** Left: AlphaFold 3 modeling of an additional interface in the Syt1 C2B–SNARE complex, involving K288 in Syt1 and E55 in SNAP25. This secondary interface recapitulates the arrangement observed in the Syt1–SNARE crystal structure (ref). Right: AlphaFold 3 modeling of the SNT-1 C2B–SNARE complex, identifying a similar secondary interface mediated by conserved K308 in SNT-1 and E55 in RIC-4. R281 (Syt1) and R301 (SNT-1), belonging to the primary interface, are also shown as reference points. **(B)** A distinct SNT-1–SNARE modeling (different from the one in Figure 1), highlighting R217 and R251 residues in the SNT-1 C2A domain and their predicted binding sites in RIC-4 (T140, N141). This modeling does not reproduce the tertiary interface seen in the Syt1– SNARE crystal structure, where D218 in syntaxin and D57 in synaptobrevin engage R199 and R233 in the Syt1 C2A domain. **(C, D)** Example traces and quantification of evoked EPSCs recorded from *snt-1* mutants rescued by wild-type SNT-1, SNT-1^K308A^, SNT-1^R217A/R251A^, and SNT-1^K308A/R217A/R251A^ (in 1 mM Ca²⁺ conditions). Mutations in these residues did not affect evoked EPSCs (amplitude or charge transfer). **(E, F)** Representative traces of mEPSCs and mIPSCs recorded in 1 mM Ca²⁺ conditions, along with quantification of their frequencies and amplitudes for the same genotypes. These mutations did not affect Ca²⁺-dependent spontaneous release. **(G, H)** Representative traces and quantification of Ca²⁺-independent spontaneous release (recorded in 0 mM Ca²⁺). Frequencies of mEPSCs and mIPSCs were significantly reduced by the R217A/R251A mutations and by the triple R217A/R251A/K308A mutations. These findings demonstrate that these residues in SNT-1 are critical for Ca²⁺-independent neurotransmitter release, likely through potential interactions with the SNARE complex. Data are presented as mean ± SEM (** *p* < 0.01, *** *p* < 0.001 vs. SNT-1 rescue; n.s., not significant; one-way ANOVA).

To assess the functional significance of residues corresponding to those in the secondary and tertiary interfaces of the Syt1–SNARE complex, we first mutated K308 in SNT-1 (K308A) to disrupt the predicted secondary interface. This mutation had no effect on either evoked or spontaneous neurotransmitter release under Ca²⁺-containing conditions (Figure 4C–F), suggesting that the secondary interface is not essential for Ca²⁺-dependent synaptic transmission. We next introduced R217A/R251A double mutations and K308A/R217A/R251A triple mutations in SNT-1, targeting residues analogous to those forming the tertiary interface or both secondary and tertiary interfaces in Syt1. Similar to the K308A mutation, these substitutions did not alter evoked or spontaneous release in the presence of Ca²⁺, indicating that these potential SNARE-contacting residues are not required for Ca²⁺-triggered SV fusion. Consistently, in SNT-3, the analogous mutations—K159A (secondary interface), and R73A/R107A (tertiary interface)—also did not affect evoked release (Figure 4—Supplemental 1).

In contrast, under 0 mM Ca²⁺ conditions, both the R217A/R251A and K308A/R217A/R251A mutations in SNT-1 significantly reduced the frequencies of mEPSCs and mIPSCs (Figure 4G, H). These results suggest that additional SNARE interactions—possibly mediated by the C2A domain—play a specific role in Ca²⁺-independent spontaneous neurotransmitter release. Together, our findings support a model in which SNT-1 engages multiple SNARE-interacting surfaces to regulate distinct modes of synaptic exocytosis, with different structural determinants contributing to Ca²⁺-dependent versus Ca²⁺-independent release.

### Synaptic transmission is abolished by disrupting all potential SNT-1–SNARE interfaces

Given the substantial preservation of mEPSCs and mIPSCs in the SNT-1 quintuple mutant under 1 mM Ca²⁺ conditions (Figure 2), we next investigated whether Ca²⁺-dependent spontaneous release could persist when all predicted SNARE-binding interfaces of SNT-1 are simultaneously disrupted. To this end, we generated an SNT-1 construct carrying the five mutations targeting the primary C2B–SNARE interface, along with additional mutations disrupting the secondary (K308A) and tertiary (R217A/R251A) interfaces—referred to as SNT-1^Octuple^—and expressed it in *snt-1* null mutants.

As expected, SNT-1-dependent evoked EPSCs were completely abolished in animals expressing the SNT-1^Octuple^ mutant, resembling the phenotype of *snt-1* null mutants (Figure 5A, B). Strikingly, spontaneous release was also nearly eliminated, with both mEPSC and mIPSC frequencies reduced to the *snt-1* mutant levels under both Ca²⁺-containing and Ca²⁺- free conditions (Figure 5C–F). These findings indicate that while the primary C2B–SNARE interface is essential, additional SNARE contacts—likely mediated by both the C2A and C2B domains—are also required for spontaneous SV fusion. Together, these results support a model in which SNT-1 interacts with the SNARE complex through multiple, partially redundant interfaces. This multivalent binding strategy likely ensures robust synaptic transmission across a range of physiological conditions.

**Figure 5.**
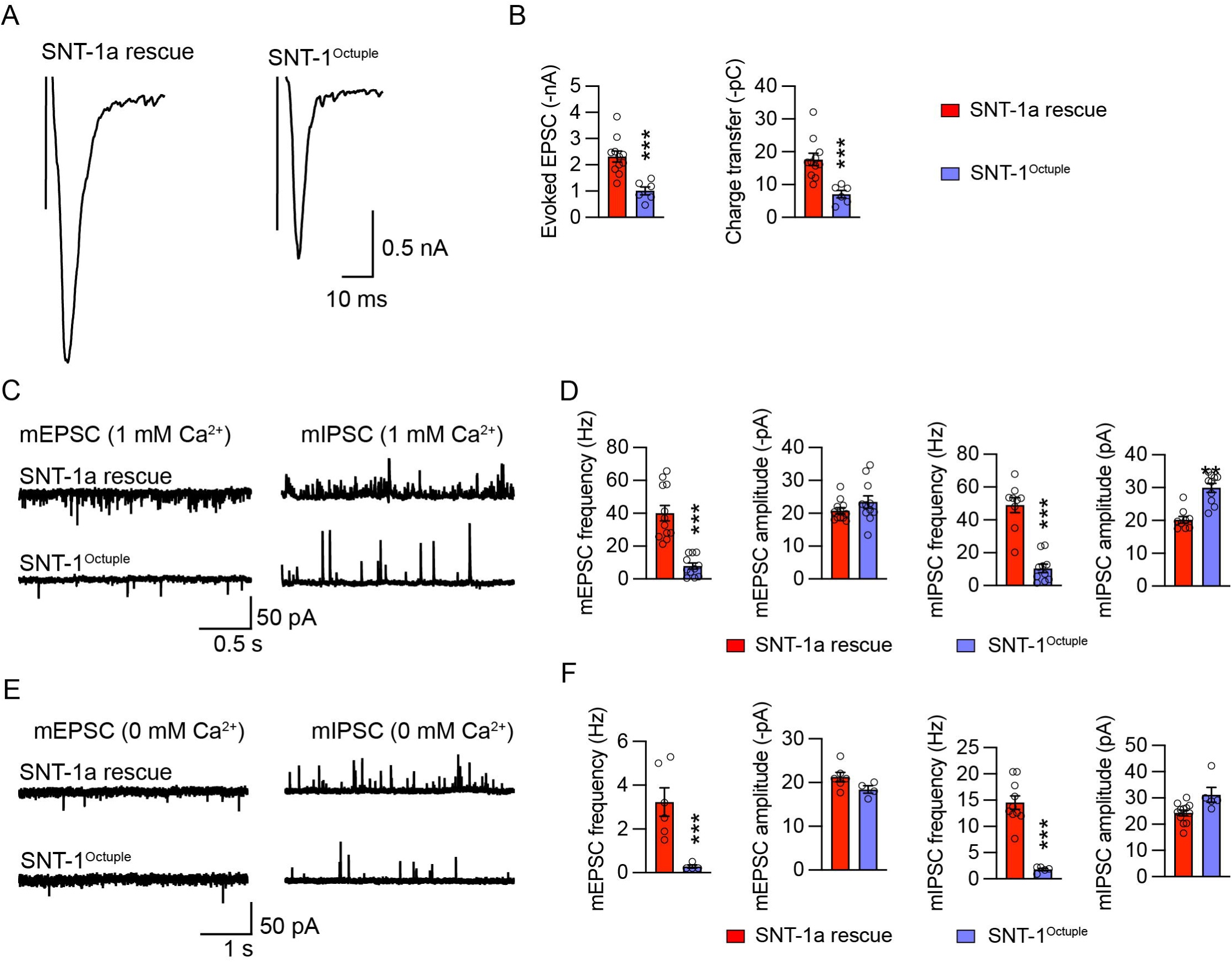
Mutations of SNT-1 residues corresponding to all primary, secondary, and tertiary interfaces in the Syt1-SNARE eliminate synaptic transmission. **(A,B)** Evoked EPSCs recorded from *snt-1* mutants rescued by wild-type SNT-1 and SNT-1^Octuple^. The octuple mutations include the quintuple mutations, K308A, R217A, and R251A. Like the quintuple mutations, the octuple mutations completely abolished evoked EPSCs. **(C-F)** Representative traces and quantifications of mEPSCs and mIPSCs recorded under 1mM Ca^2+^ conditions (C, D) and 0mM Ca^2+^ conditions (E, F) from wild-type SNT-1 and SNT-1^Octuple^ rescued animals. The octuple mutations nearly abolished both Ca^2+^-dependent and Ca^2+^-independent spontaneous release. Data are presented as mean ± SEM (** *p* < 0.01, *** *p* < 0.001 vs. SNT-1 rescue; one-way ANOVA).

### The polybasic motifs are critical for SNT-1 and SNT-3 function

In addition to SNARE binding, the C2 domains of synaptotagmins—including Syt1—play essential roles in SV exocytosis by interacting with the plasma membrane. These interactions are mediated through two mechanisms: Ca²⁺ binding by the C2 domain loops—via five aspartate residues in each domain, and electrostatic interactions involving positively charged polybasic motifs. Both the C2A and C2B domains of Syt1 contain Ca²⁺-binding loops that insert into the plasma membrane upon Ca²⁺ binding (Bai et al., 2000; Bai et al., 2004; Bradberry et al., 2019; Chapman and Davis, 1998; Grushin et al., 2019). However, only Ca²⁺ binding to the C2B domain is essential for triggering fast, synchronous neurotransmitter release (Fernandez-Chacon et al., 2002; Lee et al., 2013; Nishiki and Augustine, 2004a). Similar mechanisms have been reported for *C. elegans* SNT-1 and SNT-3 (Li et al., 2021; Li et al., 2018a).

The polybasic motifs are thought to mediate electrostatic interactions with the negatively charged inner leaflet of the plasma membrane (Bai et al., 2004; Bradberry et al., 2019; Loewen et al., 2006; Park et al., 2015). Sequence alignment shows that these motifs are highly conserved in both SNT-1 and SNT-3 (K207/K208/K209/K210 in SNT-1 C2A, K344/K345/K346/K347/K351 in SNT-1 C2B; K63/K64/ K65/K66 in SNT-3 C2A, K196/K197/K198/K202 in SNT-3 C2B; Figure 6A, B; Figure 6-Supplemental 1A, B; Figure 1-Supplemental 3;), suggesting they play conserved and essential roles in synaptic exocytosis. Structurally, these residues form a relatively flat, positively charged surface on the outer face of the C2B domain (Figure 6A, B), likely facilitating efficient membrane association. Based on AlphaFold models and structural analyses, we propose a working model in which the C2B domain engages the plasma membrane through this polybasic surface while simultaneously interacting with SNAREs via distinct binding sites (Figure 6C). This arrangement may promote rapid insertion of Ca²⁺-binding loops during synaptic activity.

**Figure 6.**
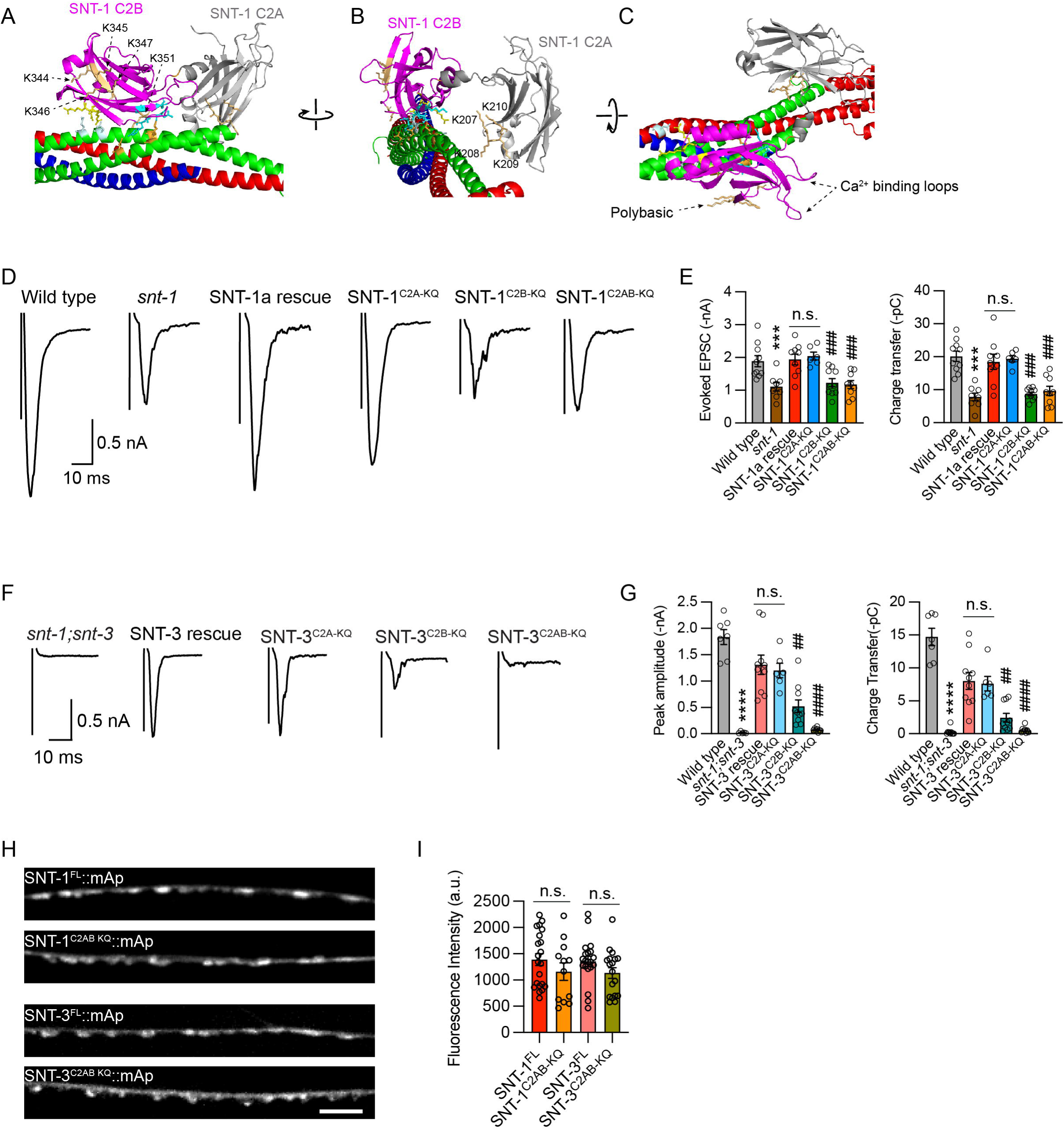
Effects of the polybasic motifs in the C2 domains of SNT-1 and SNT-3 on evoked fast and slow neurotransmitter release. **(A,B)** Positions of the polybasic motifs in the C2A (K207/K208/K209/K210) and C2B (K344/K345/K346/K347/K351) domains of SNT-1, shown in front (A) and side (B) views. **(C)** A proposed working model of the SNT-1 C2B domain, in which C2B interacts with the plasma membrane through its flat polybasic patch and with the SNARE complex via the primary interface. **(D, E)** Evoked EPSC traces (D) and quantification of amplitude and charge transfer (E) from *snt-1* mutants rescued with wild-type SNT-1 and SNT-1 carrying polybasic motif mutations in the C2A domain (SNT-1^C2A-KQ^), C2B domain (SNT-1^C2B-KQ^), or both domains (SNT-1^C2AB-KQ^). Polybasic mutations in the C2B domain, but not in the C2A domain, abolished evoked fast release. **(F, G)** Evoked EPSC traces (F) and quantification (G) from *snt-1;snt-3* double mutants rescued with wild-type SNT-3 or SNT-3 carrying polybasic mutations in C2A (SNT-3^C2A-KQ^), C2B (SNT-3^C2B-KQ^), or both domains (SNT-3^C2AB-KQ^). In SNT-3, both C2A and C2B polybasic motifs are required for evoked slow release. **(H, I)** Representative confocal z stack images for mApple-tagged SNT-1^FL^, SNT-1^C2AB-KQ^, SNT-3^FL^, and SNT-3^C2AB-KQ^ (all driven by the *unc-129* promoter), and the quantification of the fluorescence intensity from the lines. Scale bar, 5 μm. Data are presented as mean ± SEM (** *p* < 0.01, *** *p* < 0.001 vs. wild-type; ### *p* < 0.001 vs. *snt-1* mutant; n.s., not significant compared to SNT-1 rescue; one-way ANOVA).

To test the functional relevance of these polybasic motifs in fast and slow neurotransmitter release, we generated charge-neutralizing lysine-to-glutamine mutations (K→Q) in either the C2A, C2B, or both C2 domains of SNT-1 and SNT-3. These mutant variants—SNT-1^C2A-KQ^, SNT-1^C2B-KQ^, SNT-1^C2AB-KQ^, SNT-3^C2A-KQ^, SNT-3^C2B-KQ^, SNT-3^C2AB-KQ^—were then tested for their ability to rescue evoked EPSCs in *snt-1* and *snt-1;snt-3* mutant backgrounds. In SNT-1, mutation of the C2B polybasic motif nearly abolished the SNT-1-dependent fast evoked release, whereas mutation of the C2A motif had no significant effect (Figure 6D, E). A similar pattern was observed for SNT-3: disruption of the C2B motif impaired release, while C2A mutations alone did not. However, unlike SNT-1, a substantial level of evoked EPSCs remained in SNT-3^C2B-KQ^ rescued animals. Only when both C2 domains were mutated (SNT-3^C2AB-KQ^) was evoked release eliminated (Figure 6F, G).

These results demonstrate that polybasic motifs are essential components of the C2 domains in both SNT-1 and SNT-3, but they contribute differently to synaptic exocytosis. The stronger dependence of SNT-1 on the C2B polybasic motif is consistent with its role as a fast-release Ca²⁺ sensor, whereas the combined contributions of both C2 domains in SNT-3 may reflect the mechanistic diversity underlying slow-release triggering. These findings highlight distinct molecular strategies by which fast and slow Ca²⁺ sensors mediate SV fusion.

We next investigated whether the polybasic mutations affect the expression levels of SNT-1 and SNT-3, potentially contributing to the observed changes in SV release. To test this, we generated mApple-tagged fusion proteins of wild-type and mutant SNT-1 and SNT-3 (SNT-1^FL^::mApple, SNT-1^C2AB-KQ^::mApple, SNT-3^FL^::mApple, and SNT-3^C2AB-KQ^::mApple), all driven by the *unc-129* promoter, and examined their abundance in the nervous system. Quantitative fluorescence analysis showed that the polybasic mutations in both C2 domains did not affect the expression levels of either SNT-1 or SNT-3, as fluorescence intensities were indistinguishable between control and mutant strains (Figure 6H, I). These results indicate that the SV release defects caused by the polybasic mutations are due to functional impairments, rather than altered protein expression. This conclusion is consistent with previous findings in cultured cortical neurons (Wu et al., 2022).

### Polybasic motifs contribute to both Ca²⁺-dependent and -independent spontaneous release

We next examined how mutations of the polybasic motifs influence spontaneous neurotransmitter release. Under 1 mM Ca²⁺ conditions, the mEPSC frequency was significantly reduced by approximately 50% in animals expressing SNT-1 with polybasic mutations in the C2B domain, whereas mutations in the C2A domain had no significant effect (Figure 7A, B). This pattern mirrors the effects of the SNARE-binding mutations of the primary interface (Figure 2), emphasizing the essential role of the C2B domain in mediating Ca²⁺-dependent spontaneous release. In contrast, the frequency mIPSC frequency was not altered by polybasic mutations in either the C2A, C2B, or both domains, suggesting that distinct mechanisms contribute to SNT-1–mediated Ca²⁺-dependent GABA release (Figure 7C, D). One possibility is that SNARE interactions—potentially through the primary, secondary, and tertiary interfaces—play a more dominant role in inhibitory motor neurons, compensating for the loss of polybasic motif-mediated membrane interactions.

**Figure 7.**
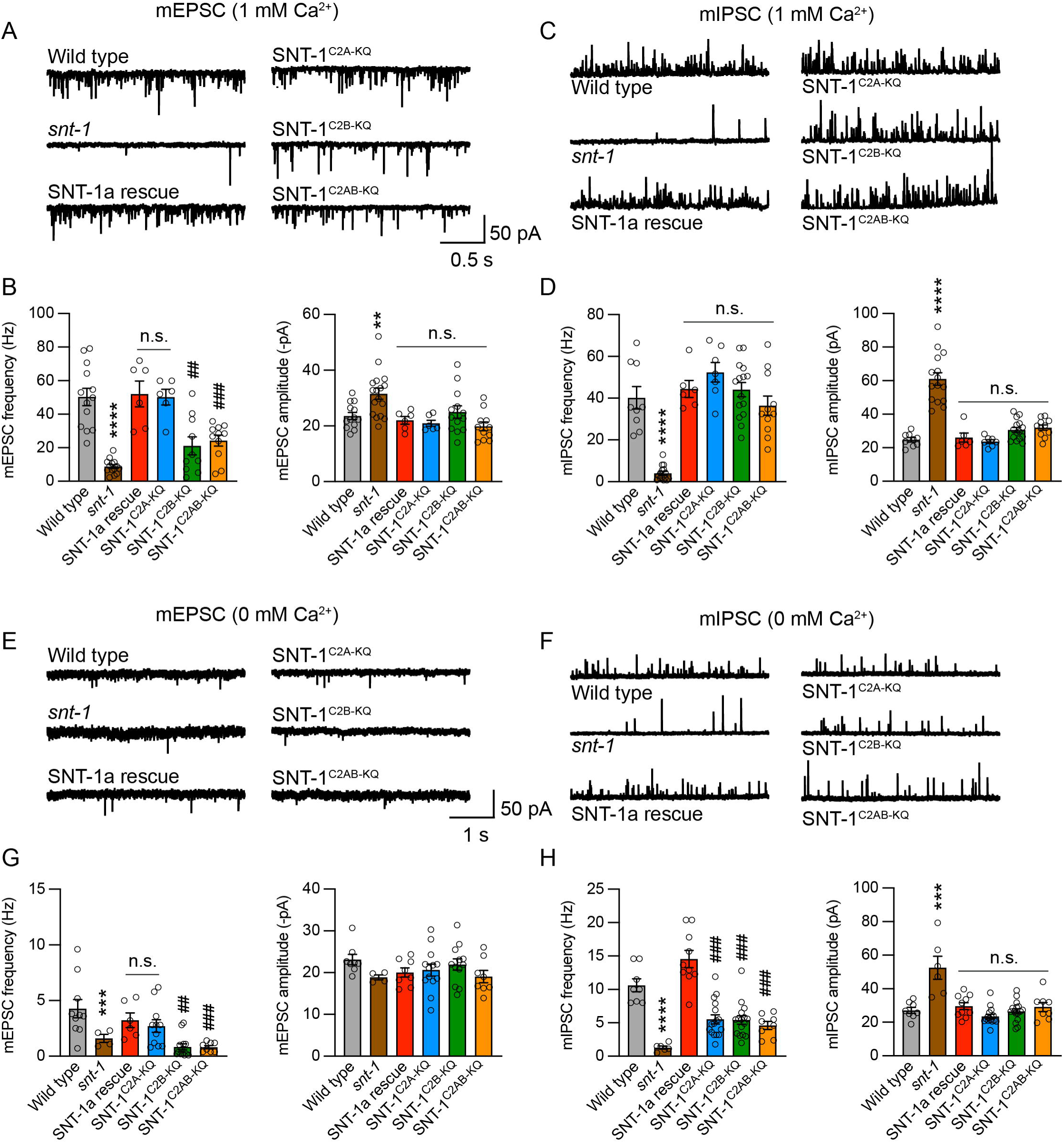
The polybasic motifs in SNT-1 are required for spontaneous neurotransmitter release. **(A-D)** mEPSCs and mIPSCs recorded from wild-type, *snt-1* mutant, and *snt-1* mutant rescued with SNT-1 containing polybasic mutations in C2A, C2B, or both C2 domains, under 1 mM Ca²⁺ conditions. The polybasic motif in the C2B domain plays a dominant role in supporting mEPSCs, without affecting mIPSCs. **(E-H)** Representative traces and quantification of Ca²⁺-independent mEPSCs and mIPSCs recorded under 0 mM Ca²⁺ conditions. In the absence of Ca²⁺, both polybasic motifs in C2A and C2B domains contribute similarly to mIPSCs, whereas only the polybasic motif in the C2B domain regulates mEPSCs. Data are presented as mean ± SEM (** *p* < 0.01, *** *p* < 0.001 vs. wild-type; ### *p* < 0.001 vs. *snt-1* mutant; n.s., not significant compared to SNT-1 rescue; one-way ANOVA).

We also assessed Ca²⁺-independent spontaneous release by recording mEPSCs and mIPSCs under 0 mM Ca²⁺ conditions. Notably, both mEPSC and mIPSC frequencies were markedly reduced in animals rescued with the SNT-1^C2B-KQ^ variant (Figure 7E-H), indicating a critical role for the C2B polybasic motif in Ca²⁺-independent neurotransmitter release as well. Interestingly, a similar reduction in mIPSC frequency was observed in SNT-1^C2A-KQ^– rescued animals, implicating the C2A domain in Ca²⁺-independent GABA release.

These findings are further supported by the observation that Ca²⁺-independent mIPSCs were also reduced in animals expressing the SNT-1^R217A/R251A^ mutant (Figure 4), pointing to a cooperative function of distinct regions within the C2A domain in mediating Ca²⁺-independent release. Structural analysis of the Syt1 and SNT-1 C2A domains reveals that the polybasic residues form a flat, positively charged surface located on the same face as R217 and R251 (Figure 7-Supplemental 1), although the precise orientation of the C2A domain during SV fusion remains unclear. Nonetheless, this spatial arrangement and the functional analyses suggest that the polybasic patch and the conserved arginine residues in SNT-1 C2A may act cooperatively to facilitate Ca²⁺-independent neurotransmitter release.

## Discussion

Following our previous work identifying SNT-1 and SNT-3 as the primary and secondary Ca²⁺ sensors that trigger fast and slow neurotransmitter release, respectively, in this study we uncovered both conserved and divergent mechanisms by which the dual Ca²⁺ sensors in *C. elegans* regulate synaptic exocytosis through their C2 domains. Focusing on the residues involved in the Syt1–SNARE primary interface, we found that the C2B–SNARE interaction plays a highly conserved role in both SNT-1-mediated fast release and SNT-3-mediated slow release. Our results also demonstrated that the polybasic motifs in SNT-1 and SNT-3 are critical for evoked fast and slow neurotransmitter release. These findings align with studies in rodents, supporting the idea of evolutionarily conserved mechanisms for synaptic transmission mediated by synaptotagmin. Importantly, our results also revealed distinct mechanisms by which SNT-1 and SNT-3 trigger fast and slow release, including differential contributions of the two subregions within the C2B-SNARE primary interface, as well as of the polybasic motifs in the C2A and C2B domains. Finally, our findings uncovered previously unknown roles of SNT-1 in supporting spontaneous release. Below, we discuss the significance of these findings.

### C2B–SNARE primary interface is conserved from invertebrates to vertebrates

Biochemical data from prior studies have demonstrated interactions between Syt1 and the SNARE complex (Chapman et al., 1995; Chapman and Jahn, 1994; Sutton et al., 1995; Wang et al., 2016; Zhang et al., 2002). However, only relatively recently has the crystal structure of the Syt1–SNARE complex revealed precise binding sites between Syt1 and SNAREs (Zhou et al., 2015). The two binding regions, located on opposite ends of the C2B domain, form a large and stable contact, ensuring the C2B domain play a critical role for SV fusion. The AlphaFold 3 modeling, together with our functional analyses, confirms that this C2B–SNARE primary interface is conserved from invertebrates to vertebrates (in SNT-1, SNT-3 and Syt1) and is critical for the function of these Ca²⁺ sensors in synaptic transmission, including evoked fast and slow release. The pan-neuronal expression of SNT-1 and SNT-3 (Li et al., 2021) suggest that the C2B-SNARE primary interaction in this dual Ca^2+^ sensors system trigger neurotransmitter release in all neurons in *C. elegans*.

In fact, sequence alignment shows that the residues involved in the Syt1–SNARE primary interface are conserved in only three of the seven synaptotagmin isoforms in *C. elegans*: SNT-1, SNT-2, and SNT-3. However, SNT-2 is primarily expressed in the intestine, where it acts as a Ca²⁺ sensor for neuropeptide secretion (Wang et al., 2013). Because the SNARE proteins that mediate membrane fusion in the intestine likely differ from those in motor neurons, it remains unclear how SNT-2 triggers vesicle fusion in the intestine. Indeed, AlphaFold 3 models of SNT-2 with the motor neuron SNAREs (UNC-64, RIC-4, and SNB-1) do not show a similar C2B binding pattern as observed in SNT-1 and SNT-3. These findings suggest that the C2B–SNARE primary interface may be a unique characteristic of synaptotagmins involved in neuronal vesicle fusion.

Despite the structural information of Syt1-SNARE, SNT-1-SNARE, and SNT-3-SNARE, the mechanistic actions of the C2 domains during fusion remain still unclear. It has been proposed that the C2B and SNAREs form the primary interface before Ca^2+^-triggering, as these interactions occur in a Ca^2+^-independent manner (Masumoto et al., 2012; Zhou et al., 2015). Our results of the dramatic decrease in Ca^2+^-independent spontaneous release by the quintuple mutations support this notion. Despite the critical roles of the C2B-SNARE primary interaction, our results showed that the C2A domain is also indispensable for synaptic transmission, likely through cooperation with the C2B domain (see below discussion).

### The function of the C2A domain in SNT-1

The C2A domain has received less attention since the finding that disrupting its Ca^2+^ binding capacity did not cause significant changes in basal synaptic transmission in cultured hippocampal neurons (Fernandez-Chacon et al., 2002). This finding has been confirmed by later studies in *Drosophila* Syt1 (Lee et al., 2013) and *C. elegans* SNT-1 (Li et al., 2018a). In contrast, mutations that abolish the Ca^2+^-binding to C2B completely eliminate evoked synchronous release (Lee et al., 2013; Li et al., 2018a; Nishiki and Augustine, 2004a). However, the C2A domain is clearly important, as synaptic transmission is abolished by removing C2A from Syt1 or SNT-1, or replacing it with an additional C2B domain in *Drosophila* Syt1 (Lee et al., 2013; Li et al., 2018a). These findings suggest that although the C2B domain is the primary driver of release, it requires the presence of the C2A domain, which likely plays an important supportive role. However, the precise mechanism by which C2A contributes to this process remains yet to be solved.

In this study we explored functional roles of the C2A domain in SNT-1 by focusing on the conserved polybasic motif and the R217 and R251 residues—corresponding to R199 and R233 in rat Syt1—which are implicated in mediating the C2A-SNARE tertiary interface (Zhou et al., 2015). We found that mutations of these residues significantly reduced spontaneous release in the absence of Ca^2+^, suggesting they may mediate Ca^2+^-independent interactions with SNAREs or the plasma membrane. Interestingly, these residues also play critical roles in spontaneous release in the presence of Ca^2+^. Specifically, mIPSCs were completely abolished by the R217A/R251A mutations in the quintuple mutant background (Figure 5), indicating that the C2A and C2B domains have redundant roles in Ca^2+^-dependent spontaneous GABA release, highlighting the cooperation of C2A and C2B in synaptic transmission. Furthermore, we discovered that the C2A domain is involved in evoked slow neurotransmitter release mediated by SNT-3. While polybasic mutations in SNT-3 C2A alone did not affect evoked release, they abolished the residual slow release when combined with C2B polybasic mutations (Figure 6). Collectively, these findings demonstrate the functional importance of the C2A domain within the dual Ca^2+^ sensor system in *C. elegans*.

### SNT-1 regulation of spontaneous release

Although the worm SNT-1 shares similar mechanisms with mammalian Syt1 in triggering evoked fast neurotransmitter release, their regulation of spontaneous release differs—indeed, it is opposite. Spontaneous release is dramatically enhanced in Syt1 knockout neurons (Kochubey and Schneggenburger, 2011; Lee et al., 2013; Maximov and Sudhof, 2005; Xu et al., 2009; Yoshihara and Littleton, 2002), whereas it is markedly decreased in *snt-1* mutants (Li et al., 2021; Li et al., 2018a). The hypothesized model for Syt1 inhibition on spontaneous release is that it clamps the SNARE complex, thereby preventing excessive vesicle fusion under physiological conditions. However, how Syt1 integrates the two seemingly opposing functions—promotion of evoked release and inhibition of spontaneous release—remains unclear. Functional analysis by Brunger’s group revealed that these two regulatory modes can be distinguished by the E295A/Y338W mutations in region 1 of the Syt1 C2B–SNARE primary interface, providing direct evidence that the two binding regions within this interface play determinant roles in controlling Syt1’s dual functions. Recent studies by Rosenmund’s group showed that mutations in SNAP25 that are involved in region I and II of Syt1 C2B-SNARE interface cause differential changes in spontaneous release, providing additional evidence for differential roles of the two binding regions (Toulme et al., 2024). Interestingly, opposite regulation of spontaneous and evoked neurotransmitter release has also been observed in mutants lacking another SNARE-binding protein, complexin (Huntwork and Littleton, 2007; Martin et al., 2011; Maximov et al., 2009). The function of complexin is highly conserved across worms, flies, and mammals, and prior studies have shown that its distinct functional domains mediate its regulation of these two modes of neurotransmitter release. All these findings suggest that interactions with the SNARE complex by the key synaptic proteins determine how spontaneous release occurs.

Our prior work showed that SNT-1 regulation of spontaneous release requires both C2 domains. mEPSCs and mIPSCs are not changed by Ca²⁺-binding mutations in either C2A or C2B alone, but are significantly decreased by mutations in both C2 domains (Li et al., 2018a). However, these Ca^2+^-binding mutations in C2 domains do not account for the dramatic reduction of Ca^2+^-independent spontaneous release in *snt-1* mutants (Figure 2, Supplemental-1). In this study, we demonstrated that Ca²⁺-independent spontaneous release requires the C2B–SNARE primary interaction (Figure 2). These results are consistent with the Ca²⁺-independent primary interaction of Syt1 C2B–SNARE. Furthermore, we found that Ca²⁺-dependent spontaneous release also requires additional interactions mediated by residues corresponding to those in the secondary and tertiary interfaces of the Syt1–SNARE complex (Figure 5). These findings highlight the multiple mechanisms by which SNT-1 regulates spontaneous release. Notably, although both the primary and secondary interfaces of the Syt1–SNARE complex were successfully identified by AlphaFold 3, the tertiary interface—mediated by C2A—could not be predicted. Thus, it remains unclear how the two arginine residues (R217 and R251) in the C2A domain are involved in regulating spontaneous release. One possibility is that these SNT-1 residues also mediate interactions with the SNAREs, similar to their roles in Syt1 C2A, thereby contributing to spontaneous release.

Interestingly, our results also showed that, in contrast to its inhibitory function in mammals, Syt1 restored the majority of spontaneous release in *snt-1* mutants, demonstrating that Syt1 can promote spontaneous release in worms—similar to its homolog SNT-1. This suggests that the distinct roles of Syt1 and SNT-1 in regulating spontaneous release in worms and mammals are likely not intrinsic to Syt1 and SNT-1 themselves, but instead reflect differences in the underlying fusion machineries of these two systems, such as subtle variations in their SNARE complexes. Future experiments testing whether SNT-1 can suppress the high-frequency spontaneous release observed in Syt1 knockout neurons, as well as cross-species rescue experiments with SNARE proteins, would provide valuable insights into the distinct regulatory mechanisms of SNT-1 and Syt1 in spontaneous release.

### Distinct mechanisms employed by SNT-1 and SNT-3?

Although SNT-1 and SNT-3 have distinct physiological roles in triggering fast and slow neurotransmitter release, respectively, AlphaFold 3 modeling suggests that they share a similar mode of interaction with the SNARE complex via their C2B domains. Both isoforms exhibit conserved binding residues and comparable orientations of C2B relative to the SNARE bundle (Figure 1), supporting a conserved mechanism in which the C2B–SNARE unit acts as a functional entity upon Ca²⁺ influx (Zhou et al., 2015). Functional evidence supports this model: mutations in region 2 arginine residues (R418A/R419A in SNT-1 and R269A/R270A in SNT-3) disrupt evoked fast and slow release (Figure 3), highlighting the importance of region 2 in maintaining C2B–SNARE complex stability. However, our results with the EY mutations (E315A/Y358W in SNT-1 and E166A/Y209W in SNT-3) suggest that the mechanisms by which SNT-1 and SNT-3 trigger release may not be entirely identical. Despite similar structural interfaces, the impact of these mutations differs between the two isoforms, pointing to potential intrinsic differences in their function.

Several explanations may account for this discrepancy. In region 1 of the Syt1 C2B–SNARE interface, four key residues (E295, K297, N336, Y338 in rat Syt1) are conserved in both SNT-1 and SNT-3. In the EY mutant background, the remaining K and N residues may still interact with the SNAP-25 homolog RIC-4 (specifically residues E37 and N161, Figure 1), allowing partial preservation of the region 1 interface and overall C2B–SNARE structure. This residual interaction may suffice to maintain slow release (SNT-3), but fall below the threshold required for the more rapid and synchronous release mediated by SNT-1. In contrast, region 2 contains only three conserved arginine residues (R281, R398, R399 in Syt1), and mutating two of them (R398A/R399A)—or their corresponding residues in SNT-1 and SNT-3—destabilizes the C2B–SNARE interaction entirely, abolishing both fast and slow release. These findings suggest that while both SNT-1 and SNT-3 employ a shared core mechanism via the C2B–SNARE interface, they may differ in their sensitivity to partial disruption, possibly due to additional structural or regulatory differences unique to each isoform.

The polybasic motifs may also contribute to the distinct mechanisms of SNT-1 and SNT-3. In both Syt1 and SNT-1, the C2B domain contains four consecutive lysine residues (K344/K345/K346/K347 in SNT-1), along with two nearby lysines (K351/K352 in SNT-1). By contrast, the SNT-3 C2B domain has only three consecutive lysines (K196/K197/K198) and two adjacent ones (K204/K206) (Figure 1 Supplemental-3). This difference may allow the polybasic motif in SNT-1 C2B to anchor more strongly to membranes than that in SNT-3 (Figure 6). In addition, the presence of a neighboring glutamate residue (E195) near the polybasic motif in SNT-3 may further weaken its membrane association through electrostatic repulsion. These distinctions likely underlie the differential roles of SNT-1 and SNT-3 in mediating fast versus slow neurotransmitter release. Notably, Syt7—the secondary Ca²⁺ sensor for slow, asynchronous release in mammals—features a nearly identical polybasic motif to SNT-3 (K319/K320/K321 and K325/K326), along with the same adjacent glutamate (E318) (Figure 6 Supplemental-2), suggesting this configuration may be a conserved feature of slow Ca²⁺ sensors.

Our data also highlight the contribution of the C2A domain to the distinct mechanisms of SNT-1 and SNT-3, as evidenced by the effects of polybasic motif mutations. A proposed model is that the polybasic motif within the C2B domain anchors it near the plasma membrane, facilitating rapid insertion of the Ca²⁺-binding loops upon Ca²⁺ influx (Figure 6A, Figure 6 Supplemental-1). However, differences in the orientation or positioning of the C2A domain between SNT-1 and SNT-3 may modulate their membrane-binding properties, thereby influencing their distinct kinetic roles in fast and slow neurotransmitter release.

Our findings in this study also raised several intriguing questions about the dual Ca²⁺ sensor system in *C. elegans*. For instance, do SNT-1 and SNT-3 compete for SNARE complex binding *in vivo*? Would swapping their C2 domains result in a switch in their functional identity? Could SNT-3 function as a fast Ca²⁺ sensor if targeted to the membrane via a transmembrane domain? Addressing these questions in future studies will further deepen our understanding of the mechanistic divergence between SNT-1 and SNT-3 and may shed light on how synaptic transmission has evolved across species.

## Materials and Methods

### RESOURCE AVAILABILITY

#### Lead Contact

Further information and requests for resources and reagents should be directed to and will be fulfilled by the Lead Contact, Zhitao Hu.

#### Materials Availability

All new strains and plasmids created in this study will be provided on request with no restrictions.

#### Data And Code Availability

This study did not generate/analyze any code.

### EXPERIMENTAL MODEL AND SUBJECT DETAILS

Strain maintenance and genetic manipulation were performed as previously described (Brenner, 1974). Animals were cultivated at room temperature on nematode growth medium (NGM) agar plates seeded with OP50 bacteria. On the day before experiments L4 larval stage animals were transferred to fresh plates seeded with OP50 bacteria for all the electrophysiological recordings. The following strains were used:

Wild-type, N2 bristol

NM204 *snt-1(md290)*

CX1034 *snt-3(ky1034)*

ZTH795 *snt-1(md290); snt-3(ky1034)*

ZTH403 *hztEx001 [Psnb-1::SNT-1]; snt-1(md290)*

ZTH389 *hztEx389 [Psnb-1::Syt1]; snt-1(md290)*

ZTH1284 *hztEx451 [Psnb-1::SNT-1^Quintuple^]; snt-1(md290)*

ZTH698 *hztEx149 [Prab-3::SNT-3]; snt-1(md290); snt-3(tm5776)*

ZTH1149 *hztEx438 [Prab-3::SNT-3^Quintuple^]; snt-1(md290); snt-3(ky1034)*

ZTH1053 *hztEx419 [Psnb-1::SNT-1^E315A/Y358W^]; snt-1(md290)*

ZTH1026 *hztEx397 [Psnb-1::SNT-1^R418A/R419A^]; snt-1(md290)*

ZTH1140 *hztEx431 [Prab-3:a:SNT-3^E166A/Y209W^]; snt-1(md290); snt-3(ky1034)*

ZTH1172 *hztEx441 [Prab-3::SNT-3^R269A/R270A^]; snt-1(md290); snt-3(ky1034)*

ZTH1052 *hztEx417 [Psnb-1::SNT-1^K308A^]; snt-1(md290)*

*Z*TH1050 *hztEx415 [Psnb-1::SNT-1^R217A/R251A^]; snt-1(md290)*

ZTH1220 *hztEx445 [Psnb-1::SNT-1^R217A/R251A/K308A^]; snt-1(md290)*

ZTH1143 *hztEx433 [Prab-3:SNT-3^R73A/R107A^]; snt-1(md290); snt-3(ky1034)*

ZTH1134 *hztEx425 [Prab-3:SNT-3^K159A^]; snt-1(md290); snt-3(ky1034)*

ZTH1381 *hztEx455 [Psnb-1::SNT-1^Octuple^]; snt-1(md290)*

ZTH1265 *hztEx447 [Psnb-1::SNT-1^C2A-KQ^]; snt-1(md290)*

ZTH1041 *hztEx399 [Psnb-1::SNT-1^C2B-KQ^]; snt-1(md290)*

ZTH1273 *hztEx449 [Psnb-1::SNT-1^C2AB-KQ^]; snt-1(md290)*

ZTH1405 *hztEx460 [Prab-3::SNT-3^C2A-KQ^]; snt-1(md290); snt-3(ky1034)*

ZTH1064 *hztEx421 [Prab-3::SNT-3^C2B-KQ^]; snt-1(md290); snt-3(ky1034)*

ZTH1066 *hztEx423 [Prab-3::SNT-3^C2AB-KQ^]; snt-1(md290); snt-3(ky1034)*

ZTH394 *hztEx380 [Punc-129::SNT-1::mApple]; NuIs165 [Punc-129::UNC-10::GFP]*

ZTH1432 *hztEx464 [Punc-129::SNT-1^C2AB-KQ^::mApple]; NuIs165 [Punc-129::UNC-10::GFP]*

ZTH673 *hztEx381 [Punc-129::SNT-3::mApple]; NuIs165 [Punc-129::UNC-10::GFP]*

ZTH1436 *hztEx468 [Punc-129::SNT-3^C2AB-KQ^::mApple]; NuIs165 [Punc-129::UNC-10::GFP]*

### METHOD DETAILS

#### Constructs, transgenes and germline transformation

The *snb-1* promoter (3kb) and *unc-129* promoter (2.6kb) were inserted into JB6 vector between the SphI and BamHI sites, all the SNT-1 and SNT-3 expression constructs were amplified by PCR and inserted into JB6 vector between the KpnI and NotI sites. Red fluorescent protein mApple was inserted between NotI and MluI sites in-frame with *snt-1* or *snt-3* genes for imaging experiments.

Transgenic strains were isolated by microinjection of various plasmids using Pmyo-2::NLS-mCherry (KP#1480) as the co-injection marker.

#### Behavioral assay

The thrashing behavior of *C. elegans* was measured to assess their locomotion ability. A single thrash is defined as a complete cycle in which the worm’s body bends to its maximum angle and then returns to its initial position. For the assay, worms at the D1 stage were washed with M9 buffer and gently transferred to NGM agar plates containing 1 mL of M9 buffer without OP50 bacteria. In each trial, 10-15 worms were selected and allowed to acclimate in the M9 buffer for 1 minute before recording. They were then observed swimming freely in the buffer for 2 minutes and 30 seconds. Worm identification and thrashing behavior were analyzed using the WormLab System. Only individuals exhibiting detectable thrashing activity for more than 30 seconds were considered valid samples. Approximately 30 samples were analyzed in each experimental group.

#### Electrophysiology

Electrophysiology was conducted on dissected *C. elegans* as previously described (Li et al., 2019; Liu et al., 2019). Worms were superfused in an extracellular solution containing 127 mM NaCl, 5 mM KCl, 26 mM NaHCO_3_, 1.25 mM NaH_2_PO_4_, 20 mM glucose, 1 mM CaCl_2_, and 4 mM MgCl_2_, bubbled with 5% CO_2_, 95% O_2_ at 22°C. The 1 mM CaCl_2_ was replaced by 1 mM MgCl_2_ to record mEPSCs and mIPSCs in 0 mM of Ca^2+^. Whole-cell recordings were carried out at -60mV for all EPSCs, including mEPSCs, evoked EPSCs, and sucrose-evoked responses. The holding potential was switched to 0mV to record mIPSCs. The internal solution contained 105 mM CH_3_O_3_SCs, 10 mM CsCl, 15 mM CsF, 4mM MgCl_2_, 5mM EGTA, 0.25mM CaCl_2_, 10 mM HEPES, and 4mM Na_2_ATP, adjusted to pH 7.2 using CsOH. Stimulus-evoked EPSCs were stimulated by placing a borosilicate pipette (5–10 μm) near the ventral nerve cord (one muscle distance from the recording pipette) and applying a 0.4 ms, 85 μA square pulse using a stimulus current generator (WPI).

#### Fluorescence imaging

Animals were immobilized on 2% agarose pads with 30 mM levamisole. Fluorescence imaging was performed on a spinning-disk confocal system (3i Yokogawa W1 SDC) controlled by Slidebook 6.0 software. Animals were imaged with an Olympus 100x 1.4 NA Plan-Apochromat objective. Z series of optical sections were acquired at 0.13 μm steps. Images were deconvolved with Huygens Professional version 16.10 (Scientific Volume Imaging, The Netherlands) and then processed to yield maximum intensity projections using ImageJ 1.54h (Wayne Rasband, National Institutes of Health) (Schneider et al., 2012).

### QUANTIFICATION AND STATISTICAL ANALYSIS

#### Data acquisition and analysis

All electrophysiological data were obtained using a HEKA EPC10 double amplifier (HEKA Elektronik) filtered at 2 kHz, and analyzed with open-source scripts developed by Eugene Mosharov (http://sulzerlab.org/Quanta_Analysis_8_20.ipf) in Igor Pro 7 (Wavemetrics). All imaging data were analyzed in ImageJ software. Each set of data represents the mean ± SEM of an indicated number (n) of animals. To analyze mEPSCs and mIPSCs, a 4pA peak threshold was preset, above which release events are clearly distinguished from background noise. The analyzed results were re-checked by eye to ensure that the release events were accurately selected.

#### Statistical analysis

All data were statistically analyzed in Prism 9 software. Normality distribution of the data was determined by the D’Agostino-Pearson normality test. When the data followed a normal distribution, an unpaired student’s t-test (two-tailed) or one-way ANOVA was used to evaluate the statistical significance. In other cases, a Mann-Whitney test or one-way ANOVA following Kruskal-Wallis test was used. A summary of all electrophysiological data is provided in Supplementary File 1, with the results presented as mean ± SEM.

## Acknowledgements

We thank the *C. elegans* Genetics Stock Center for strains and reagents. We thank members of the Hu lab. This work was supported by a National Institutes of Health research grant (R56NS128048 to J.R. and Z.H.), a CityU startup fund (9610647 to Z.H.), and a National Health and Medical Research Council Project grant (APP1122351 to Z.H.).

---

**Figure 1 Supplemental-1. Rescue of spontaneous release in *snt-1* mutants by rat Syt1.**

**(A, C)** Representative traces of mEPSCs (A) and mIPSCs (C) recorded from wild-type, *snt-1* mutant, SNT-1 rescue, and Syt1 rescue worms.

**(B, D)** Quantification of mEPSC (B) and mIPSC (D) frequencies and amplitudes from the same genotypes shown in (A) and (C). Data are presented as mean ± SEM (*** *p* < 0.001 vs. wild-type; # *p* < 0.05, ### *p* < 0.001; one-way ANOVA).

**Figure 1 Supplemental-2. AlphaFold 3 modeling of the rat Syt1–SNARE complex.**

**(A)** Typical AlphaFold 3 model of the Syt1–SNARE complex, showing the two binding regions (region 1 and region 2) highlighted with dashed squares. The SNARE proteins are colored red (syntaxin), green (SNAP-25), and blue (synaptobrevin).

**(B, C)** Close-up views of residues involved in region 1 (E295/K297/N336/Y338 in Syt1 and E37/K40/N159/D166 in SNAP-25) and region 2 (R281/R398/R399 in Syt1 and D51/E52/E55 in SNAP-25).

**(D)** Comparison table of residues forming the primary interface of the Syt1–SNARE, SNT-1– SNARE, and SNT-3–SNARE complexes, as determined by the real crystal structure (Zhou et al., 2015, *Nature*) and AlphaFold 3 modeling. These conserved interactions suggest a shared mechanistic role of synaptotagmins across species in SNARE-mediated synaptic vesicle fusion.

**Figure 1 Supplemental-3. Sequence alignment of Syt1, SNT-1, and SNT-3 C2 domains.**

The key residues such as the Ca^2+^ binding sites in both C2 domains, the region 1 and region 2 in the C2B domain, the polybasic sequences in both C2 domains, are indicated.

**Figure 2 Supplemental-1. The Ca^2+^-binding mutations in SNT-1 do not affect Ca^2+^-independent spontaneous release.**

**(A, C, E, G)** Example traces of mEPSCs and mIPSCs from wild type, *snt-1* mutants, and rescued strains by SNT-1, SNT-1^C2A-D3,4N^, SNT-1^C2B-D3,4N^, SNT-1^C2AB-D3,4N^.

**(B, D, F, H)** Quantification of mEPSC and mIPSC frequency and amplitude from the indicated genotypes.

Data are presented as mean ± SEM (** *p* < 0.01, *** *p* < 0.001 vs. wild type; ## *p* < 0.01 vs. SNT-1a rescue; n.s., not significant compared to SNT-1 rescue; one-way ANOVA).

**Figure 4 Supplemental-1. SNT-3 mutations of residues corresponding to Syt1–SNARE secondary and tertiary interfaces do not affect synaptic transmission.**

**(A, B)** Evoked EPSCs recorded from *snt-1;snt-3* double mutants rescued by wild-type SNT-3, SNT-3^K159A^, and SNT-1^R73A/R107A^. AlphaFold 3 modeling does not predict interactions of these SNT-3 residues with the SNARE complex, and consistent with this, the mutations did not affect evoked EPSC amplitude or charge transfer.

**(C, D)** Quantification of mEPSC and mIPSC frequencies and amplitudes for the genotypes shown in A. These mutations did not affect the spontaneous release, compared to wild-type SNT-3 rescue.

Data are presented as mean ± SEM (n.s., not significant compared to SNT-3 rescue; one-way ANOVA).

**Figure 6 Supplemental-1. AlphaFold 3 modeling of SNT-3-SNARE complex shows the position of the polybasic motifs in the C2 domains.**

(A, B) The polybasic motifs in C2B (K196, K197, K198, K202) and C2A (K63, K64, K65, K66) have similar positions as those in SNT-1 (Figure XX). (C) A potential working orientation of the C2B domain and the SNARE complex. In this model, the polybasic motif in C2A does not interact with the plasma membrane.

**Figure 6 Supplemental-2. The polybasic sequences in the C2 domains of Syt1, Syt7 and SNT-3.** Unlike the fast Ca^2+^ sensor Syt1 which contains four consecutive lysine residues in the C2B domain, the slow Ca^2+^ sensors Syt7 and SNT-3 contain three consecutive lysine residues (K196/K197/K198 in SNT-3, and K319/K320/K321 in Syt7) and one glutamate residue (E195 in SNT-3 and E318 in Syt7) in their C2B domains.

