## Supplementary figures and images for "Evolutionarily Conserved and Divergent Mechanisms of Dual Ca^2+^ Sensors in Synaptic Vesicle Exocytosis"

### Figure 1

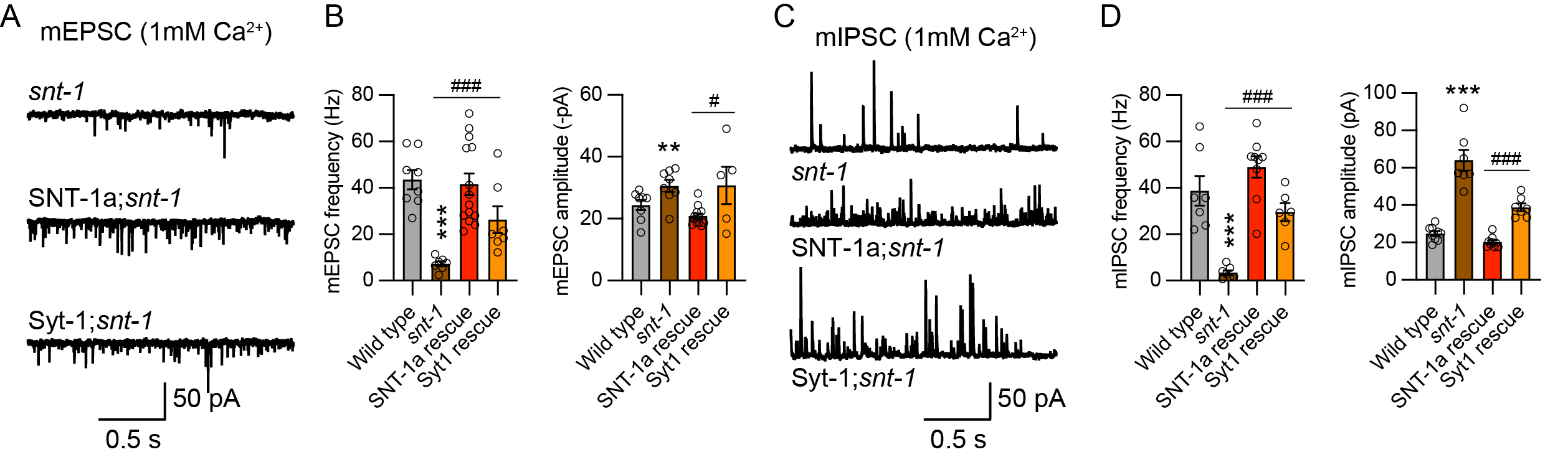

### Figure 1

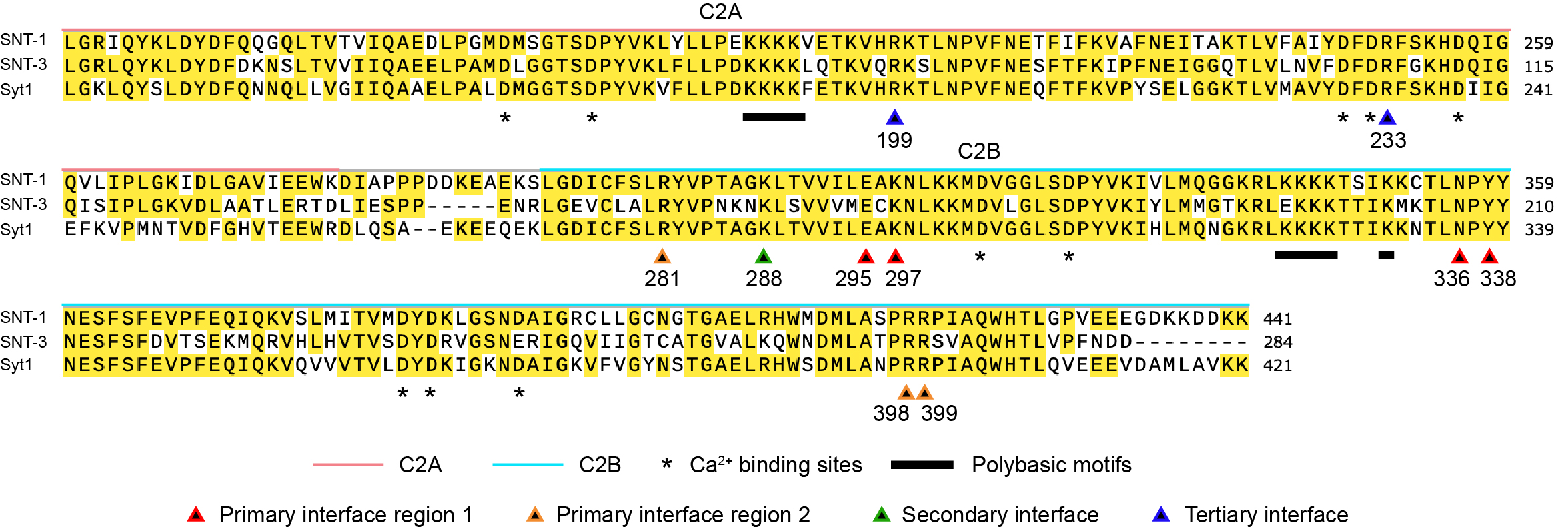

### Figure 1

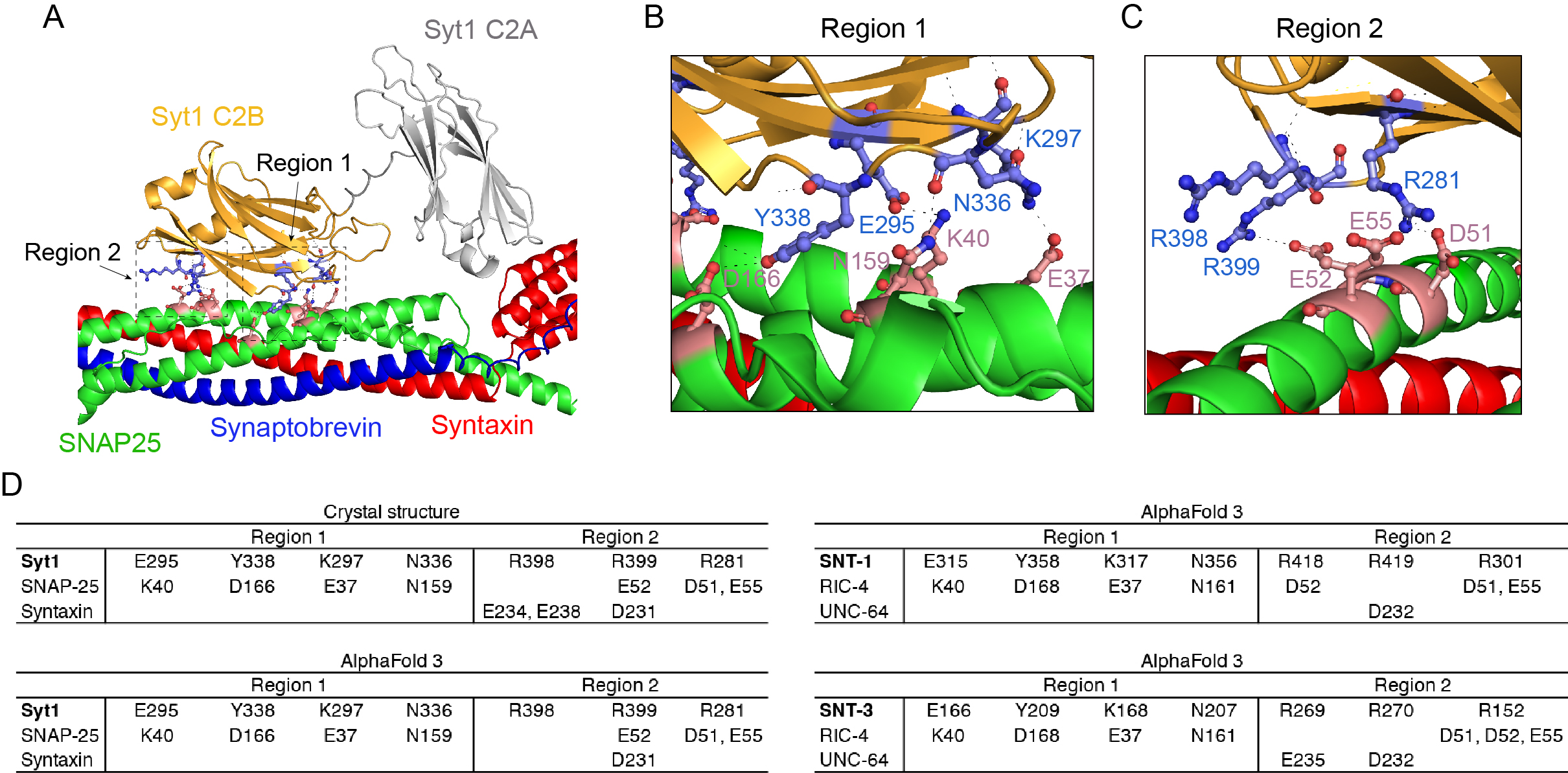

### Figure 2

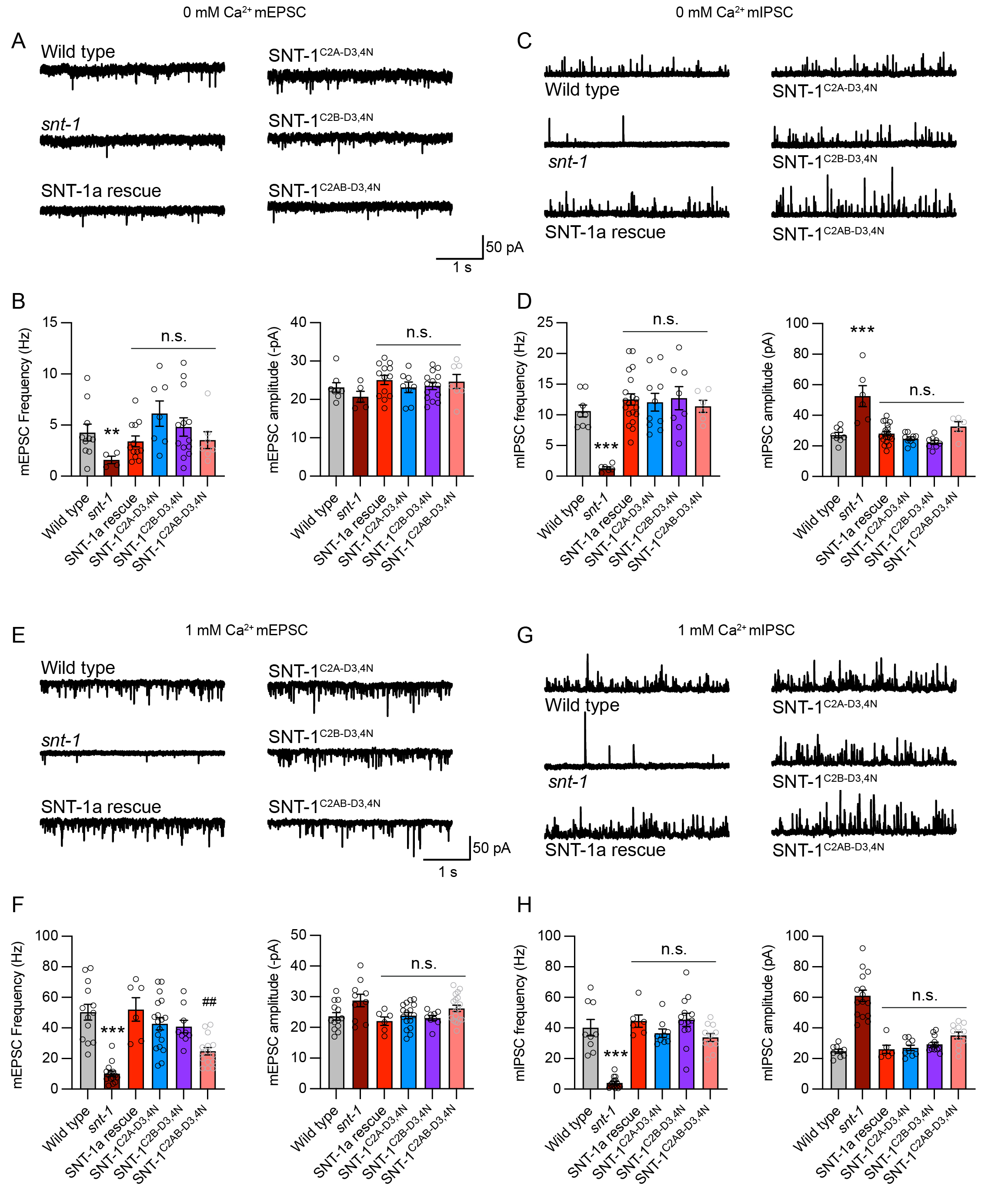

### Figure 4

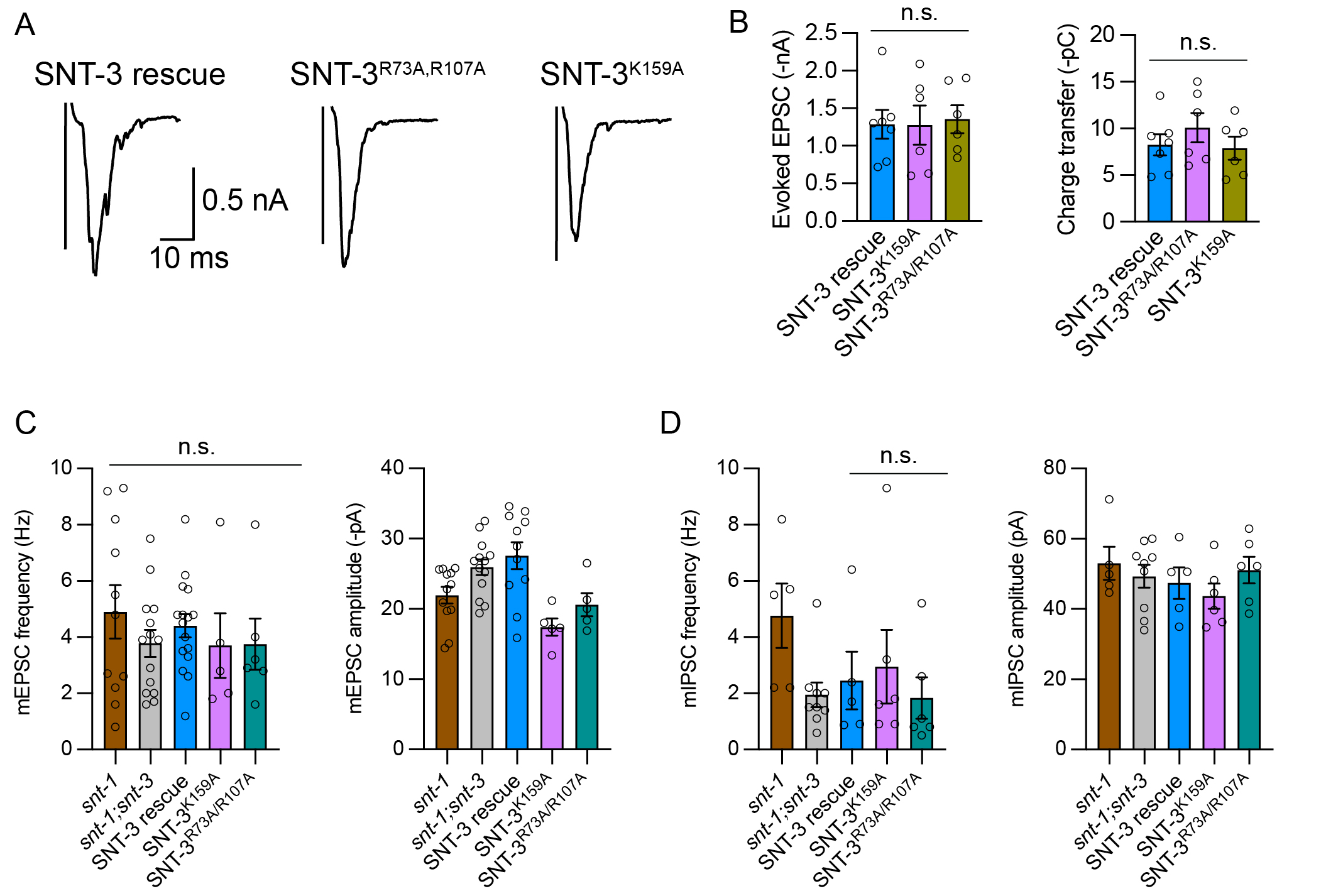

### Figure 6

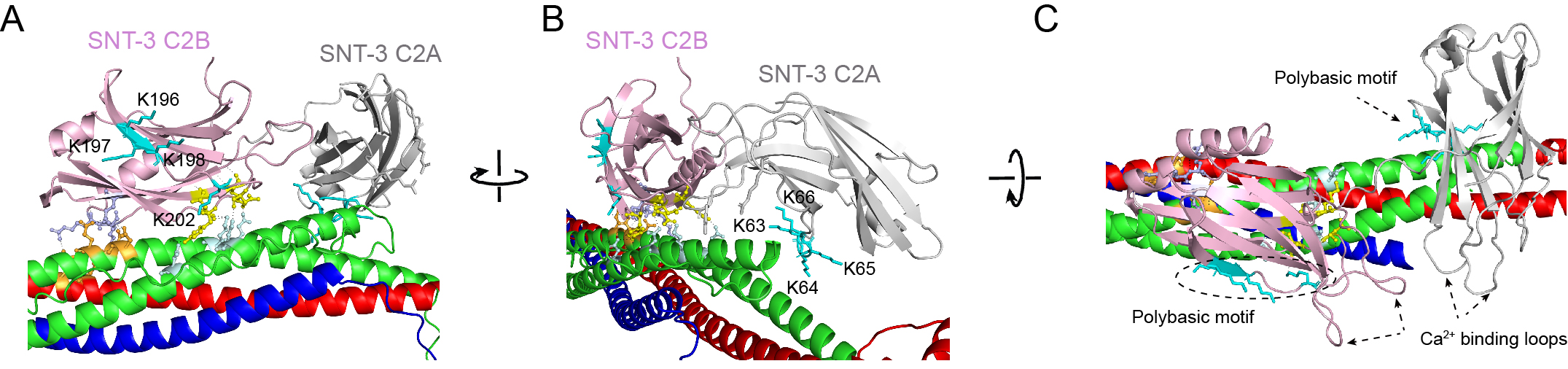

### Figure 6

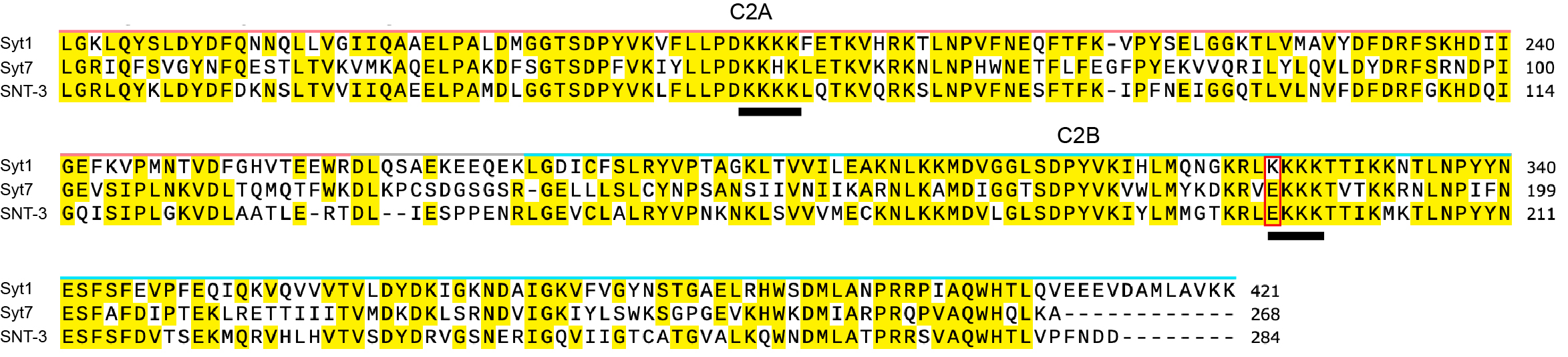

### Figure 7

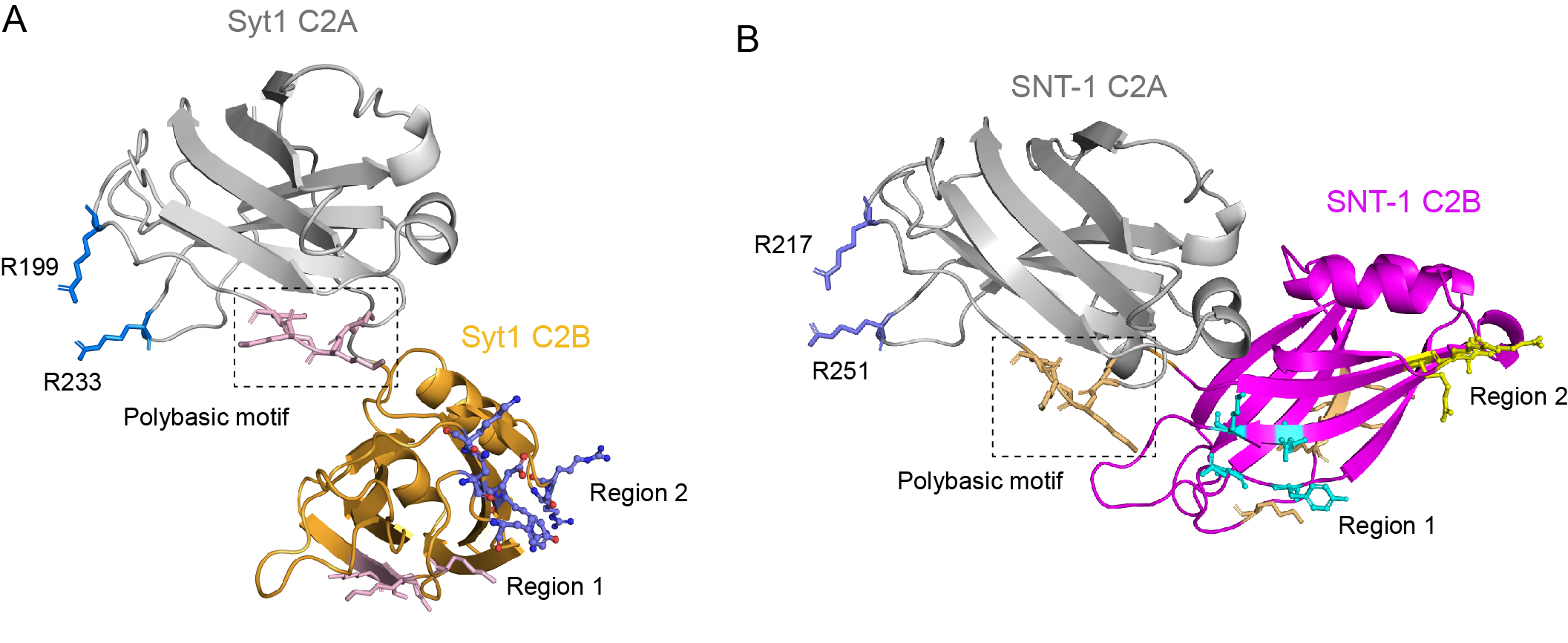
